# Oligomeric amyloid beta prevents myelination in a clusterin-dependent manner

**DOI:** 10.1101/2020.03.06.981373

**Authors:** Rebecca M. Beiter, Hannah E. Ennerfelt, Courtney Rivet-Noor, Andrea Merchak, Robin Bai, David M. Johanson, Erica Slogar, Katia Sol-Church, Jennifer L. Phillips, Tim Bathe, Christopher C. Overall, John R. Lukens, Stefan Prokop, Alban Gaultier

## Abstract

White matter loss has been described as a common occurrence in Alzheimer’s disease (AD) patients for multiple decades. However, it remains unclear why oligodendrocyte progenitor cells (OPCs) fail to repair myelin deficits in these patients. Here, we show that clusterin, a risk factor for late-onset AD, is produced by OPCs and inhibits their differentiation into oligodendrocytes. Specifically, we demonstrate that a unique subset of OPCs produces clusterin. We show that phagocytosis of debris, including amyloid beta (Aβ) and myelin, drives the upregulation of clusterin in OPCs. We confirm, *in vivo*, that Aβ oligomers drive clusterin upregulation and that OPCs phagocytose Aβ. Furthermore, we show that clusterin is a potent inhibitor of OPC differentiation and prevents the production of myelin proteins. Finally, we demonstrate that clusterin inhibits OPC differentiation by significantly reducing the production of IL-9 by OPCs. Our data reveals that clusterin may be responsible for the lack of myelin repair observed in AD and is a promising therapeutic target for AD-associated cognitive decline.

## Main

Alzheimer’s Disease (AD) is a neurodegenerative disease that currently has no approved therapeutic to effectively prevent disease progression, and patients inevitably succumb to debilitating dementia^1^. White matter loss has been documented in AD patients, but how this myelin loss contributes to disease progression remains unclear^2^. Recent studies indicate that generation of new myelin-forming oligodendrocytes is critical for memory consolidation and recall^3, 4^. Additionally, remyelination therapeutics have been shown to reduce cognitive deficits seen in an animal model of AD^5^. This evidence supports the hypothesis that the inability to generate oligodendrocytes is a significant driver of the cognitive deficits observed in AD.

The adult brain contains oligodendrocyte progenitor cells (OPCs), a highly proliferative population of glia that are maintained in a progenitor state throughout adulthood and are normally capable of generating nascent oligodendrocytes^6, 7^. However, it is not understood why OPCs fail to repair myelin damage present in AD. The ability of OPCs to produce mature, myelinating oligodendrocytes during adulthood is critical, as motor learning, memory consolidation, and memory recall are all dependent on de novo production of myelin^3, 4, 8^.

Here we investigated if clusterin, a risk factor for late-onset AD, alters the function of OPCs in AD^9^. Clusterin is a multifunctional apolipoprotein that is upregulated in multiple neurodegenerative disorders and is known to play a role in the clearance of debris^10–13^. We found that a specific subset of OPCs in the adult mouse brain express clusterin, mimicking data from humans indicating that a subset of OPCs express high levels of clusterin^14^. This same data indicates that OPCs, but not any other brain-resident cell type, upregulates clusterin expression in the context of AD^14^. Consequently, we investigated what factors might contribute to clusterin production in OPCs. We found that phagocytosis of debris, including both oligomeric Aβ and myelin debris, results in an upregulation of clusterin production by OPCs. We further discovered that clusterin is a potent inhibitor of OPC differentiation and the production of myelin proteins. At a mechanistic level, we found that clusterin reduces the production of the cytokine IL-9 by OPCs and that restoration of IL-9 levels rescues the ability of OPCs to differentiate in the presence of clusterin.

## Results

### Clusterin is expressed by a subset of OPCs

Clusterin, most commonly found as a secreted protein, is upregulated in the brains of patients with Alzheimer’s disease (Fig. 1a-b)^1^. Interestingly, we found that cells expressing clusterin RNA could be found directly surrounding Aβ plaques (Fig. 1c). This observation is conserved in pre-clinical models of AD, as clusterin is also found to be upregulated in multiple brain regions of the 5xFAD mouse model of AD (Fig. 1d, e). Because of the emerging role of OPCs and myelin in AD pathology, we investigated whether OPCs expressed clusterin in AD^2, 3^. Surprisingly, we found that OPCs express clusterin both in the AD brain, as well as in normal aging (Fig. 1f). In order to further profile these clusterin-expressing OPCs, we performed single cell sequencing on OPCs from the brains of adult mice. To accomplish this, we used an inducible *PDGFRα-CreER*; *R26-EYFP* reporter mouse to efficiently label the majority (93.56% ± 1.98) of OPCs in the brain (Extended Data Fig. 1a-d)^4^. Following the induction of YFP in *PDGFRα*+ cells, we isolated YFP^+^ cells using Fluorescence-Activated Cell Sorting (FACS) and sequenced this population (Extended Data Fig. 2a, b). We found two distinct populations of OPCs (OPC1, OPC2) in the adult brain that were observed over 3 sequencing runs and were not sex-specific (Fig. 1g, Extended Data 2c-e). Each population of OPCs expressed canonical markers and showed a unique transcriptional signature distinct from the gene expression in every other cluster (Extended Data Fig. 3a, b). Subsequent subclustering of each OPC population identified genes linked with potential unique functions for each subcluster of OPCs, and confirmed the presence of a previously identified population of proliferating OPCs (Extended Data Fig. 4a-d)^5^.

**Figure 1:**
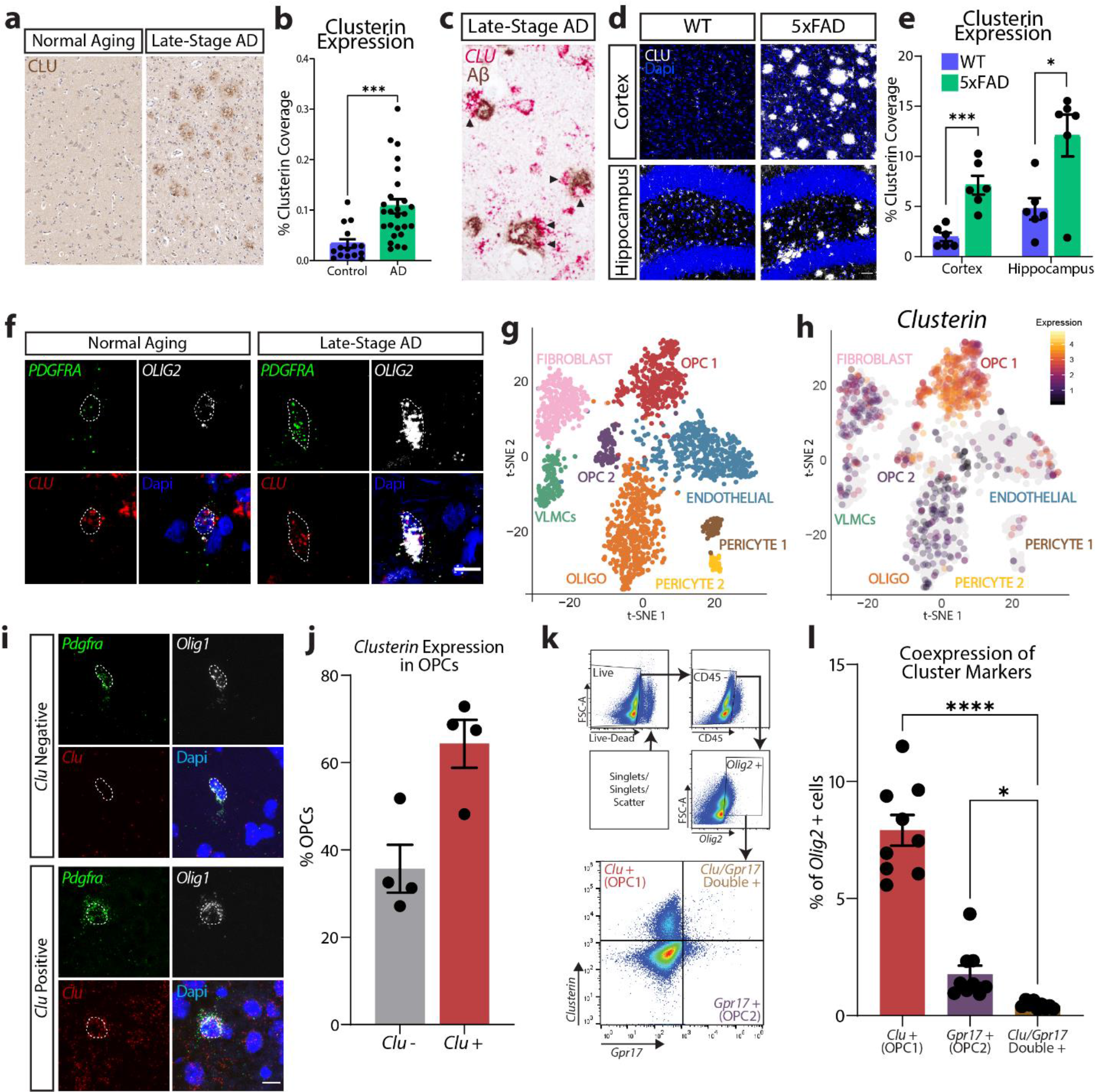
A subset of OPCs expresses the AD-risk factor clusterin. **a**, Representative image of Clusterin expression (immunohistochemistry, brown) in the cortex from a normal aging patient and a late-stage AD patient **b**, Quantification of clusterin coverage in the cortex of normal aging (*n*=15) and AD patients (*n*=26, from two independent experiments, depicted in **a**). Data analyzed using an unpaired t-test; t(39)=3.767. **c**, Detection of clusterin RNA (in situ hybridization, red) and Aβ protein (immunohistochemistry, brown) in late-stage AD brain (late-stage AD *n*=2; from two independent experiment). Arrowheads indicate clusterin-expressing cells around plaques. **d**, Representative images of clusterin expression (white) in the cortex and hippocampus of WT and 5xFAD mice. Scale bar=30µm. **e**, Quantification of clusterin coverage in the cortex and hippocampus of WT and 5xFAD mice (depicted in **c**; WT *n*=6, 5xFAD *n*=6; from two independent experiments). Statistics calculated using an unpaired Student’s t-test. Cortex: t(10)=5.049; Hippocampus: t(10)=3.119. **f**, *In situ hybridization* for OPCs (*PDGFRA* in green, *OLIG2* in white) expressing Clusterin (*CLU*; red) in normal aging and late-stage AD brains (normal aging *n*=1, late-stage AD *n*=1; from one independent experiment). Scale bar=10µm. **g**, tSNE plot of single-cell sequencing of cells isolated from PDGFRα-CreER; R26-EYFP reporter brains (*n*=6; from 3 independent experiments). Clusters were labeled with cell-type classifications based on expression of common cell-type markers. **h**, Cell-specific expression of clusterin overlaid on single-cell sequencing tSNE map. **i**, Representative images of *in situ hybridization* for OPCs (*Pdgfra* in green, *Olig1* in white) expressing or lacking clusterin (*Clu*; red). Scale bar=10µm. **j**, Quantification of *Clu*+ and *Clu-*OPCs (depicted in **i**; *n*=4 with 206 total cells analyzed; from two independent experiments). Each sample includes quantification of OPCs from the cortex, hippocampus, corpus callosum, and cerebellum. **k**, Representative gating strategy for RNA flow cytometry. **l**, Quantification of RNA flow cytometry (depicted in **j**) for *Clu* and *Gpr17* expression in *Olig2*^+^ cells in WT mice (*n*=9; from 2 independent experiments). Statistics calculated using a one-way repeated measures analysis of variance (ANOVA) with a Tukey’s post-hoc analysis. F(2,8)=90.85. *p<0.05, ***p<0.001, ****p<0.0001. All error bars represent standard error of the mean (SEM).

One cluster of OPCs (OPC1) was delineated by high clusterin expression compared to other cell types present in our dataset (Fig. 1h). We confirmed that a subset of OPCs (64.32% ± 5.49%) expressed clusterin *in vivo* using RNAscope for OPCs (*Pdgfra* and *Olig1*) and Clusterin (Fig. 1i, j). We used *Gpr17* as a marker for the remaining subset of OPCs (OPC2) and used the same method to confirm that a subset of OPCs expressed *Gpr17 in vivo* (58.77% ± 3.19%; Extended Data Fig. 5a, b). We next asked if OPC1 and OPC2 were distinct from each other, or if cells expressed markers from both clusters at the same time. We used RNA-based flow cytometry (PrimeFlow) to demonstrate that a subset of *Olig2*^+^ cells expressed clusterin (OPC1), and a mutually exclusive population expressed *Gpr17* (OPC2), with very few OPCs expressing detectable levels of both cluster markers (Fig 1.k, l). This data confirms the presence of the two distinct populations of OPCs we observed in our single-cell sequencing. Furthermore, these results show that clusterin, a risk factor for late-onset AD, is upregulated in the parenchyma of AD patients, and that this gene is expressed by a unique subset of OPCs in the mouse brain.

### Phagocytosis of oligomeric Aβ and cellular debris results in an upregulation of clusterin expression by OPCs

Given our data showing that OPCs express the AD-risk factor clusterin, we wanted to determine if OPCs interacted with Aβ plaques and could contribute to AD pathology. Surprisingly, using 5xFAD mice, we found that OPCs were surrounding and extending their processes into Aβ plaques (Fig. 2a). Because clusterin is known to facilitate debris clearance, we next wondered if OPCs were involved in the engulfment of Aβ oligomers from the brain^6^. To test this, we injected WT mice with Aβ oligomers labeled with CypHer5e, a dye that only fluoresces when it has entered the acidic environment of the lysosome. We found that OPCs around the injection site phagocytosed Aβ within 12 hours of injection (Fig. 2b, Extended Data Fig. 6a). Surprisingly, we also found that injecting Aβ was sufficient to drive an upregulation of clusterin expression in the brain (Fig. 2c, d).

**Figure 2:**
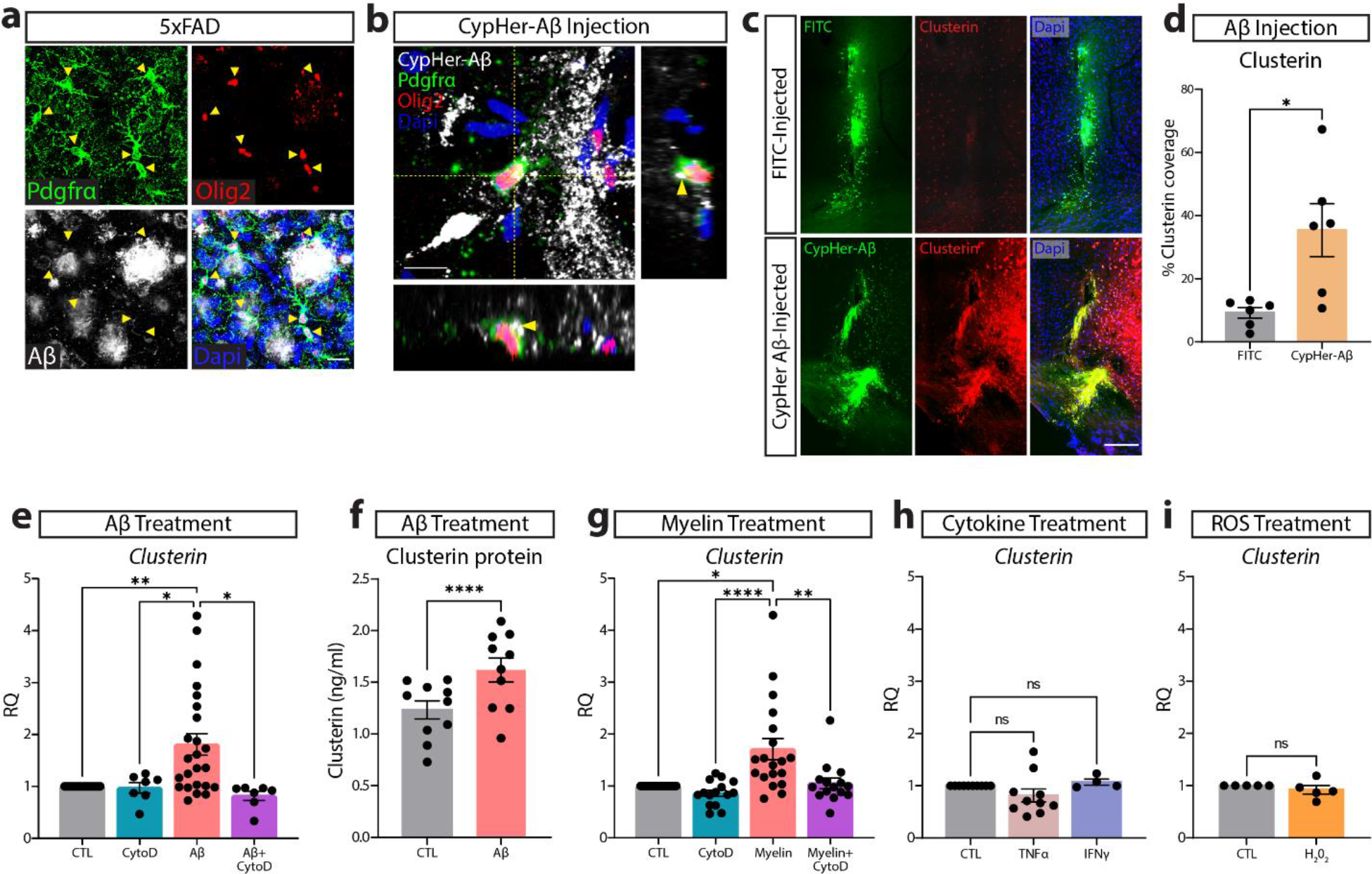
Phagocytosis of extracellular debris drives clusterin expression in OPCs. **a**, Representative image of OPCs (PDGFRα in green, Olig2 in red) surrounding Aβ plaques (white). Yellow arrowheads indicate OPCs that are extending processes into areas of Aβ accumulation (*n*=4 mice; from one independent experiment). Scale bar=20µm. **b**, Representative orthogonal view of CyPher-labeled Aβ (white) inside an OPC (PDGFRα in green, Olig2 in red) following intra-parenchymal injection of Aβ (*n*=6 mice; from one independent experiment). Yellow arrowheads indicate Aβ that can be seen inside the cell body of an OPC. Scale bar=10µm. **c**, Representative images of clusterin expression (red) in the ipsilateral (CypHer-Aβ injected; green) and contralateral (FITC injected; green) hemispheres following intra-parenchymal injection. Scale bar=100µm. **d**, Quantification of clusterin expression in the ipsilateral (Aβ-injected) and contralateral (FITC-injected) hemispheres following intra-parenchymal injection. (*n*=6 mice). Statistics calculated using a paired Student’s t-test; t(5)=2.979. **e**, qPCR analysis of clusterin expression in OPCs following a 4-hour *in vitro* treatment with 3µm Aβ and the phagocytosis blocker CytoD (1µm) or vehicle control (CTL *n*=24, CytoD *n*=7, Aβ *n*=24, Aβ+CytoD *n*=7; from seven independent experiments). Statistics calculated using a mixed effects analysis with a Tukey’s post-hoc analysis; F(1.121, 13.08) = 8.544. **f**, Quantification of clusterin protein in OPCs following 72-hour treatment with 3µM Aβ or vehicle control (CTL *n*=10, Aβ *n*=10, from two independent experiments). Statistics calculated using a paired Student’s t-test; t(9)=7.596. **g**, qPCR analysis of clusterin expression in OPCs following a 4-hour *in vitro* treatment with 100µg/ml myelin and CytoD (1µm) or vehicle control (CTL *n*=19, CytoD *n*=15, Myelin *n*=19, Myelin+CytoD *n*=15; from five independent experiments). Statistics calculated using a mixed effects analysis with a Tukey’s post-hoc analysis; F (1.389, 21.29) = 10.75. **h**, qPCR analysis of clusterin expression in OPCs following a 3-hour *in vitro* treatment with 10ng/ml TNFα or 10ng/ml IFNγ (CTL *n*=10, TNFα *n*=10, IFNγ n=4; from 2 independent TNFα experiments or 1 independent IFNγ experiment). Statistics calculated using a mixed effects analysis; F (1.106, 11.61) = 1.897. **i**, qPCR analysis of clusterin expression in OPCs following a 3-hour *in vitro* treatment with 10µm H_2_0_2_ (*n*=5; from one independent experiment). Statistics calculated using a paired Student’s t-test; t(8)=0.3417. *p<0.05, **p<0.01, ****p<0.0001, ns=not significant. All error bars represent SEM.

We next investigated whether OPCs could contribute to this Aβ-induced upregulation of clusterin. We treated primary OPCs *in vitro* with Aβ oligomers for 4 hours and observed a rapid increase in clusterin production at the RNA level (Fig. 2e). We subsequently confirmed that OPCs also upregulate the production of clusterin protein following treatment with Aβ (Fig. 2f). We found that phagocytosis of Aβ was necessary to drive this upregulation of clusterin, as treatment with cytochalasin D (CytoD), an actin polymerization inhibitor that blocks phagocytosis, prevented Aβ from increasing clusterin production in OPCs (Fig. 2e)^7–9^. Interestingly, we found that these changes in clusterin were not specific to the phagocytic clearance of Aβ oligomers, but were rather generalizable to instances in which OPCs engulfed large debris. We found that treatment of OPCs with myelin debris and apoptotic cells also produced a similar upregulation of clusterin that was abrogated by exposure to CytoD (Fig. 2g, Extended Data Figure 7). In order to eliminate the possibility that this clusterin upregulation was simply a response to any cellular stressor present in AD, we treated OPCs with other factors known to be upregulated in Alzheimer’s disease, including TNFα, IFNγ, and reactive oxygen species (ROS), which all failed to produce any change in clusterin expression (Fig. 2h, i)^10–12^.

In previous studies, clusterin has been shown to increase the phagocytic capacity of both professional and non-professional phagocytes alike^7, 8^. This led us to investigate whether clusterin altered the kinetics of phagocytosis in OPCs. Surprisingly, the addition of exogenous clusterin did not change the ability of OPCs to engulf Aβ *in vitro* (Extended Data Fig. 6b, c), indicating that phagocytosis regulates clusterin expression in OPCs, but clusterin does not subsequently regulate phagocytic ability. In sum, these results indicate that OPCs are capable of clearing extracellular debris, including Aβ, and that phagocytosis of protein or cellular debris is responsible for driving clusterin expression in OPCs.

### Exogenous clusterin inhibits OPC differentiation

The formation of new myelin has been increasingly recognized as a critical component of memory function^13, 14^. In fact, a recent study demonstrated that drugs promoting nascent myelin formation results in improved memory performance in a model of AD, indicating the importance of understanding what factors are preventing the differentiation of OPCs into oligodendrocytes^3^. A multitude of studies have demonstrated that cellular debris and protein aggregation prevent OPC differentiation, although the mechanism by which this occurs remains unclear^15, 16^. This data, along with our observation that debris clearance drives clusterin expression in OPCs (Fig. 2e, f), made us question whether clusterin might be inhibiting OPC differentiation. Consequently, we treated differentiating OPCs with exogenous clusterin and observed a striking decrease not only in genes encoding myelin proteins (*Mbp*, *Plp1*, *Cnp*), but also in *Myrf*, the master transcriptional regulator of the OPC differentiation program (Fig. 3a-d). We subsequently observed fewer MBP-positive oligodendrocytes when OPCs were differentiated in the presence of clusterin compared to clusterin-free media (Fig. 3e, f). Importantly, this decrease in differentiation is not due to OPC death, as there was no difference in OPC number following clusterin treatment (Fig. 3g). Overall, this data shows that clusterin is regulated by the phagocytosis of debris and is a potent inhibitor of OPC differentiation.

**Figure 3:**
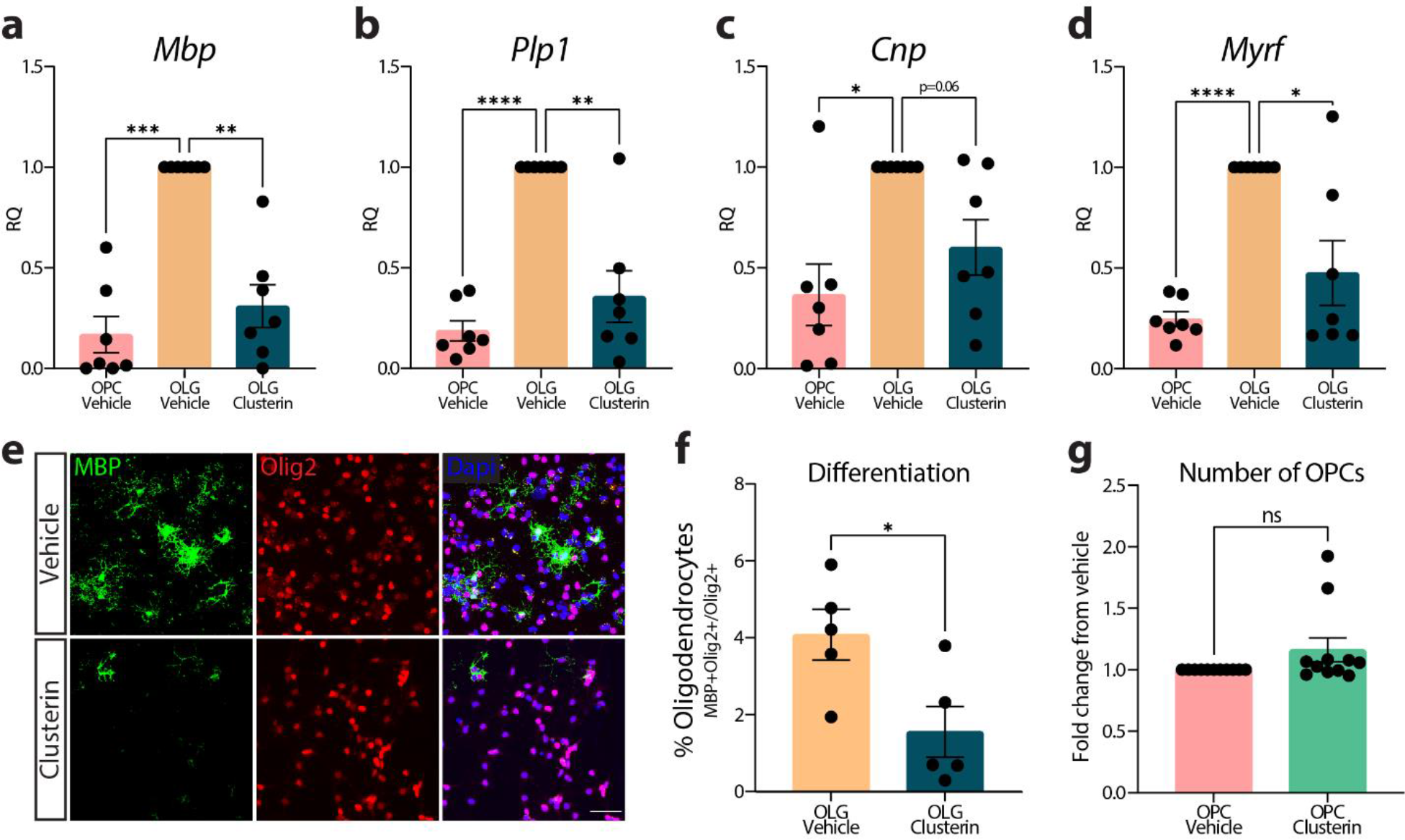
Exogenous clusterin inhibits OPC differentiation. Expression of *Mbp* (**a**), *Plp1* (**b**), *Cnp* (**c**), and *Myrf* (**d**), measured by qPCR in OPCs cultured in proliferation media (OPC Vehicle), differentiation media (OLG Vehicle), or differentiation media supplemented with 8µg/ml of clusterin (OLG Clusterin) for 72 hours (*n*=7 for all conditions; from 3 independent experiments). Statistics calculated using a repeated measures one-way ANOVA with a Tukey’s post-hoc analysis; *Mbp* F(2,6)=27.41, *Plp1* F(2,6)=25.72, *Cnp* F(2,6)=9.776, *Myrf* F(2,6)=16.41. **e**, Representative images of OPCs cultured in differentiation media with or without 8µg/ml of clusterin for 72 hours and stained for oligodendrocyte markers (MBP in green, Olig2 in red). Scale bar= 50µm. **f**, Quantification of the number of OPCs that differentiated in to oligodendrocytes following treatment with clusterin (depicted in **e**; *n*=5 for all conditions; from two independent experiments). Statistics calculated using a paired Student’s t-test; t(4)=2.780. **g**, Quantification of the number of OPCs present using a Cell Counting Kit-8 assay following a 72 hour incubation in proliferation media with or without 8µg/ml clusterin (*n*=11 for all conditions; from four independent experiments). *p<0.05, **p<0.01, ****p<0.0001, ns=not significant. All error bars represent SEM.

### Clusterin inhibits differentiation by reducing IL-9 production

We next investigated what factors might be mediating clusterin’s inhibition of OPC differentiation. OPCs have been shown to produce a variety of growth factors and cytokines that can significantly alter their local environment^17–21^. Additionally, clusterin has been shown to regulate to production of cytokines^22–24^. Based on this data, we wondered if clusterin might be inhibiting OPC differentiation by affecting growth factor and cytokine production. We performed a Luminex Assay on the supernatant of OPCs treated with vehicle or clusterin and found that clusterin altered the secretion of multiple proteins, mostly notably increasing the production of VEGF and significantly decreasing the production of IL-9 (Fig. 4a). Since growth factors from the VEGF family have been shown to induce OPC proliferation^25^, we tested whether the increase in VEGF following clusterin treatment was responsible for keeping OPCs in an undifferentiated state. However, treatment of OPCs with a combination of clusterin and an anti-VEGF antibody failed to rescue the differentiation defect observed with clusterin treatment (Extended Data Fig. 8a-c).

**Figure 4:**
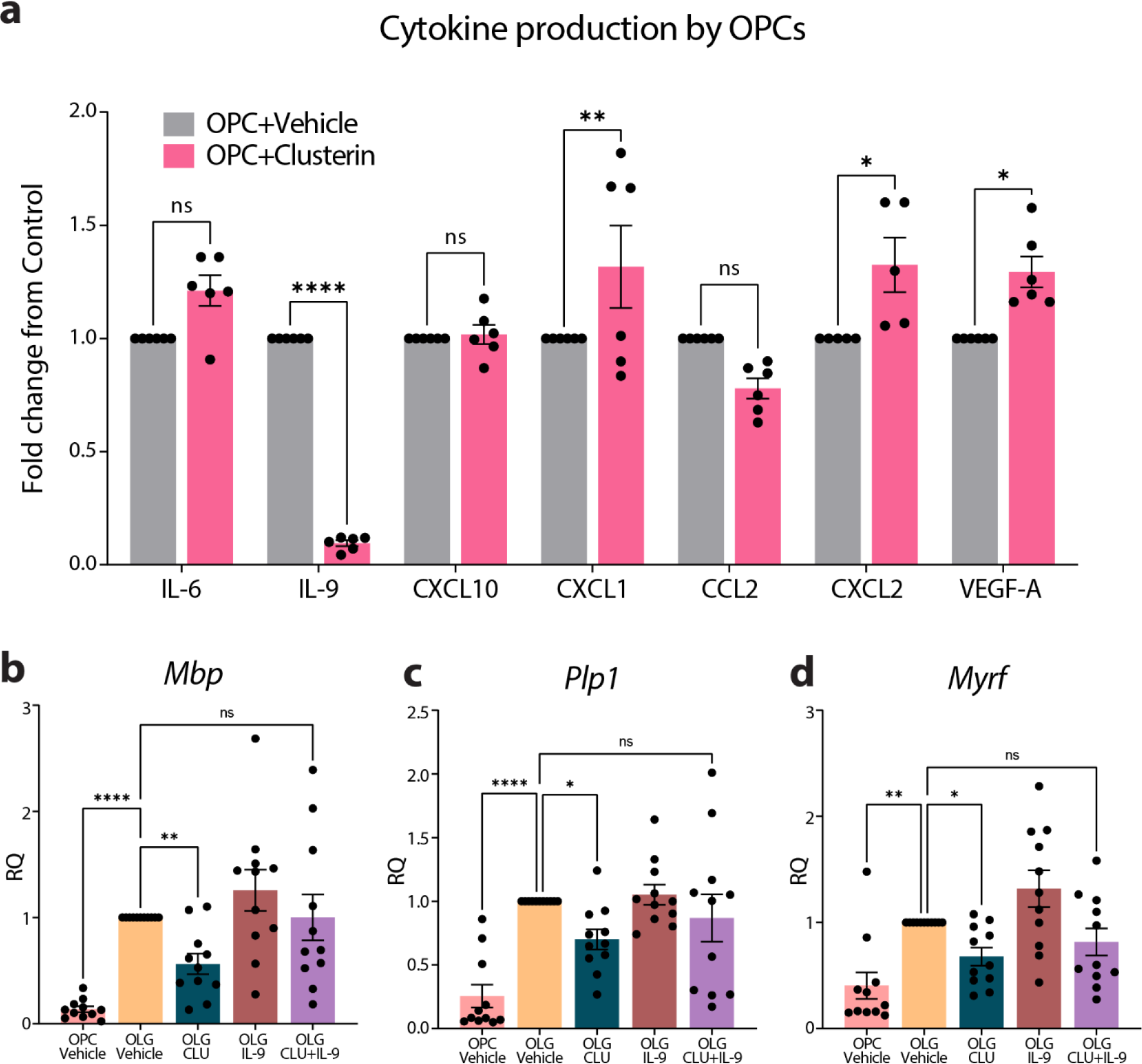
Clusterin inhibits differentiation by blocking IL-9 production. **a**, Quantification of cytokines present in the supernatant from OPCs treated with 8µg/mL clusterin or vehicle control (*n*=6 biological replicates for each condition, from two independent experiments). Data analyzed using a two-way repeated measures ANOVA with a Sidak’s multiple comparison post-hoc analysis, F (6, 34) = 23.93. Expression of *Mbp* (**b**), *Plp1* (**c**), and *Myrf* (**d**), measured by qPCR in OPCs cultured in proliferation media (OPC Vehicle), differentiation media (OLG Vehicle), differentiation media supplemented with 8µg/ml of clusterin (OLG CLU), differentiation media supplemented with 100ng/ml IL-9 (OLG IL-9), or differentiation media supplemented with 8µg/ml of clusterin and 100ng/ml IL-9 (OLG CLU+IL-9) for 72 hours (*n*=11 for all conditions; from 3 independent experiments). Statistics calculated using a repeated measures one-way ANOVA with a Tukey’s post-hoc analysis; *Mbp* F(10,40)=10.15, *Plp1* F(10,40)=11.96, *Myrf* F(10,40)=10.32. *p<0.05, **p<0.01, ****p<0.0001, ns=not significant. All error bars represent SEM.

The next candidate we assessed as the potential mechanism for the effects of clusterin on OPCs was IL-9, since its production was reduced by more than 90% following clusterin treatment (Fig. 4a). IL-9 is a relatively understudied cytokine known to be produced by T-cells^26^. Surprisingly, we found that the addition of exogenous IL-9 was sufficient to rescue the differentiation deficits induced by clusterin treatment (Fig. 4b-d). Overall, this data demonstrates that clusterin blocks the differentiation of OPCs by inhibiting the production of IL-9, and that IL-9 is an important factor in proper differentiation of OPCs.

## Discussion

Since the discovery of Aβ plaques in the brains of AD patients, therapeutic development has been focused on reducing plaque load. However, no therapeutic candidate, even when successful at reducing plaques, has succeeded in slowing disease progression^27^. Over recent years, a growing body of literature has speculated that other altered biological processes may be the driving force behind the clinical decline observed in AD patients^28^. Increasingly, OPCs and myelin have been implicated in the etiology and progression of AD symptoms. A recent study demonstrated that the ablation of senescent OPCs was sufficient to improve the memory impairment observed in the APP/PS1 model of AD^2^. Additionally, multiple studies have shown that inhibiting OPC differentiation prevents memory consolidation and recall, and that therapeutically increasing myelination can improve memory performance in AD models^3, 13, 14, 29^. Here, we offer a potential mechanism for the pervasive myelin deficits observed in Alzheimer’s disease and the memory decline associated with these deficits^30, 31^. We demonstrate that clusterin is upregulated in the brains of AD patients as well as in a mouse model of AD. We found that a specific subset of OPCs expresses clusterin under normal conditions, and that these clusterin-positive OPCs can be found in AD. Phagocytosis of debris, including Aβ oligomers, myelin, and apoptotic cells, efficiently drives the upregulation of clusterin in OPCs. Finally, we show that clusterin, through its effects on the production of IL-9, is a potent inhibitor of OPC differentiation and the production of myelin proteins.

A common single-nucleotide polymorphism (SNP) in clusterin has been recognized as a significant risk factor for late onset AD for over a decade^32^. While there are conflicting reports in the literature regarding how this SNP effects the function and accumulation of clusterin, there are multiple studies indicating that that increased levels of clusterin in the plasma of AD patients, regardless of the presence of a SNP at the CLU locus, correlates with a more rapid cognitive decline and an increase in brain atrophy^33–36^. These correlative studies demonstrating that excess levels of clusterin are associated with more severe AD symptoms are supported by studies indicating that removing clusterin in mouse models of AD results in a reduced plaque load as well as improved performance on memory tasks^37–39^. There are, however, reports in the literature indicting that clusterin might be involved in the clearance of Aβ plaques and the protection of neurons^40, 41^. These discrepancies likely indicate that clusterin is a multi-functional protein that may be beneficial when expressed at homeostatic levels, but could become detrimental when significantly upregulated in the context of disease pathology. Additionally, since there is an increasing amount of data indicating that the reduction of Aβ plaques in AD patients does not improve clinical symptoms, it is possible that clusterin’s inhibition of OPC differentiation, as described here, may be the mechanism though which increased levels of clusterin might contribute to AD progression^42^.

It remains to be determined exactly which cell types might be contributing to the increase in clusterin observed in Alzheimer’s disease. While we have demonstrated that OPCs upregulate clusterin production in response to the phagocytosis of Aβ as well as other cellular debris, and human single cell sequencing data indicates that OPCs are the only cell type in the brain that show an increase in clusterin expression in AD patients compared to healthy controls, it is known that astrocytes produce a significant amount of clusterin^43, 44^. However, while the exact cellular sources of elevated levels of clusterin in AD is still an open question, the fact that the most abundant isoform of clusterin encodes for a secreted chaperone protein indicates that clusterin from any cellular source, whether OPCs or any other cell type, can impact the function and differentiation capacity of OPCs^45^.

We were surprised to find that clusterin decreased the differentiation of OPCs by blocking the production of IL-9. While IL9 receptor expression has been documented in OPCs, this is the first time that OPCs have been shown to produce IL-9 and that IL-9 is important for the proper differentiation of OPCs^46^. While T-cells are known to be the main producers of IL-9, there is increasing evidence in the literature that OPCs can also contribute to cytokine production^17, 19, 47, 48^. The ability of OPCs to function as immunomodulatory cells is garnering increasing support in the literature, and our data indicating the importance of IL-9 in the function of OPCs offers an intriguing avenue for additional investigation^49, 50^.

It has been known that clusterin is increased in the brains of AD patients for over three decades^51^. However, the mechanism of clusterin’s effects on the progression of AD still remains unclear. With an antisense-oligonucleotide targeting clusterin having progressed to Phase 3 clinical trials for prostate cancer, our data demonstrating that clusterin negatively affects OPCs and myelin production offers a new therapeutic avenue to potentially improve myelin integrity and memory deficits plaguing AD patients^52, 53^. What’s more, our data linking clusterin and myelin dysregulation is supported by MRI data demonstrating that the clusterin risk allele is associated with a decrease in white matter integrity in healthy young adults, prior to onset of any cognitive decline^54^. Overall, our data offers a mechanism for the association between clusterin and Alzheimer’s disease pathology, and provides a foundation for the rapid repurposing of current drugs for the treatment of AD.

## Acknowledgements

The authors are supported by grants from the NINDS R41 NS110159, R56 NS120352 and R21 NS111204, and the Owens Family Foundation. D.A.R. is supported by T32 GM007055. S.M.S. is supported by T32 GM007267 and F31 NS103327, J.R.L is supported RF1 AG071996.

## Contributions

R.M.B and A.G. designed the study; R.M.B, H.E.E., C.R.N., A.M., E.S., K.S.C., J.R.L, S.P., J.L.P., T.B., and A.G. performed experiments; D.M.J., R.B. and C.C.O. performed sequencing analysis; R.M.B and A.G. analyzed and wrote the manuscript; A.G. oversaw the project.

## Methods

### Animals

*PDGFRα-CreER* mice (Jackson #018280) were crossed to *R26-EYFP* (Jackson, #006148) animals to generate *PDGFRα-CreER*; *R26-EYFP* mice, a previously described model^55^. C57BL/6J (Jackson, #000664) were purchased from Jackson or bred at the University of Virginia. 5xFAD mice (Jackson #34848) were bred at the University of Virginia and were used at 5-6 months old^56^. Mice were maintained on a 12 hour light/dark cycle with lights on at 7am. Behavior was performed on mice used in single-cell sequencing run 1. Testing consisted of sucrose preference, elevated plus maze, open field, and forced swim test. All animal experiments were approved and complied with regulations of the Institutional Animal Care and Use Committee at the University of Virginia (protocol #3918).

### Human Tissue Samples

Human post-mortem brain samples were provided by Dr. Stefan Prokop at the University of Florida’s Neuromedicine Human Brain and Tissue bank. All tissue collection and preparation was approved by the Institutional Review Board at the University of Florida. Clinical details of all human specimens analyzed are details in **Supplemental Table 1**.

### Tamoxifen injections

Tamoxifen (C8267, Sigma-Aldrich) was dissolved in corn oil at 37°C overnight at 20 mg/mL. Tamoxifen was administered *i.p*. at 200 mg/kg with a maximum dose of 4 mg per injection. For single-cell sequencing experiments, six-week old mice were given two injections of tamoxifen, three days apart. For validation of Cre recombination in *PDGFRα-CreER*; *R26-EYFP* brains, five to six week old mice were injected with 0, 1, 2, or 3 doses of tamoxifen, each given three days apart. For those mice receiving three doses of tamoxifen, the final dose was given at 150 mg/kg.

### OPC Culture

OPCs were cultured as previously described, with a few modifications^57^. Briefly, postnatal cortices (P0-P4) were rapidly dissected and meninges removed. The tissue was digested in 2ml of Accutase (Gibco, A1110501) supplemented with 50 units/mL DNase (Worthington Biochemical, LS002139). Cells were then passed through a 70µm filter and grown in suspension in neurosphere media consisting of DMEM/F12 (Gibco, 11320082), B27 (Gibco, 17504044), Pen-Strep (Gibco, 15140122), and 10ng/mL EGF (Peprotech, 315-09). Following expansion as neurospheres, cells were switched to oligosphere media consisting of DMEM/F12 (Gibco, 11320082), B27 (Gibco, 17504044), Pen-Strep (Gibco, 15140122), 10ng/mL FGF (Peprotech, 450-33), and 10ng/mL PDGF-ΑA (Peprotech, 315-17). Cells were allowed to grow in suspension for at least 2 days. Cells were then plated as attached OPCs on 0.01% Poly-L-Lysine coated plates (Electron Microscopy Sciences, 19320-B) in the same media. Cells were allowed to attach for at least 12 hours and then subsequent assays were performed.

### OPC Proliferation

OPCs were plated in proliferation media (described above) supplemented with 8µg/mL clusterin (Sino Biological 50485-M08H) or an equivalent volume of vehicle (H_2_0 or PBS) for the control samples. OPCs were allowed to proliferate for 40-72 hours. Cell number was assessed using the Cell Counting Kit-8 (Dojindo, CK04) according to manufacturer’s instructions. Optical density (OD) was measured at 450nm. For data analysis, the OD of a media only control was subtracted from the OD of all experimental samples. The OD of wells treated with clusterin were then normalized to the biologically identical control well.

### OPC Differentiation

OPCs were plated in proliferation media (described above) or differentiation media consisting of DMEM/F12 (Gibco, 11320082), B27 (Gibco, 17504044), Pen-Strep (Gibco, 15140122), 10ng/mL FGF (Peprotech, 450-33), 10ng/mL CNTF (Peprotech, 450-13), and 40ng/mL T3 (Sigma, T6397). For clusterin conditions, 8µg/mL clusterin (Sino Biological 50485-M08H) was added to the differentiation media and an equivalent volume of vehicle (H_2_0 or PBS) was added to the differentiation and proliferation control samples. For IL-9 rescue experiments, 100ng/ml IL-9 (Peprotech 219-19) was added to the differentiation media and an equivalent volume of vehicle (0.1% BSA) was added to the differentiation and proliferation control samples. For VEGF inhibition experiments, 10µg/ml of a VEGF function-blocking antibody (R&D Systems MAB9947-SP) was added to differentiation media and 10µg/ml of control IgG (Biolegend 403501) was added to the differentiation and proliferation control samples. OPCs were allowed to differentiate for 48-72 hours and were then subsequently processed for RNA extraction and qPCR or for immunofluorescence.

### Aβ Preparation and *in vitro* treatment

Aβ oligomers were prepared as previously described^58^. Briefly, human Amyloid Beta1-42 (Echelon Biosciences, 641-15) was dissolved in HFIP (Sigma, 52517) to make a 1mM solution and was allowed to desiccate overnight. The resulting peptide film was diluted to a 5mM solution in DMSO and subsequently diluted to a 100µM solution in phenol-free F-12 cell culture media (Gibco, 11039-021) and allowed to incubate overnight at 4°C. For the analysis of clusterin expression following Aβ-treatment, OPCs were treated with 3µM Aβ or a vehicle control for 4 hours. For experiments that included samples treated with CytoD, cells were pretreated for 30 minutes with 1µM CytoD (Millipore-Sigma, C8273) or DMSO and subsequently treated with 3µM Aβ or vehicle control (also containing CytoD or DMSO) for 4 hours. For CypHer-labeled Aβ experiments, 400µM Aβ oligomers in DMSO and phenol-free F-12 cell culture media was incubated with an equivalent volume of 0.1M sodium bicarbonate (Fisher Scientific, S233-500) and 400µM CypHer5e (Cytiva, PA15401) for 30 minutes at room temperature. Following incubation, the solution was spun through a buffer exchange column (Thermo Scientific, 89882) to remove any excess dye.

### Myelin Preparation and *in vitro* treatment

Myelin was prepared from mouse brains as previously described^59^. Purified myelin was passed through an insulin syringe prior to use to ensure cells were treated with a homogenous solution. Cells were treated with 100µg/ml myelin or vehicle control for 4 hours. For two replicates that included samples treated with CytoD, cells were pretreated for 30 minutes with 1µM CytoD (Millipore-Sigma, C8273) or DMSO and subsequently treated with 100µg/ml myelin or vehicle control (also containing CytoD or DMSO) for 4 hours. For the remaining two replicates that included samples treated with CytoD, no pretreatment was performed. Pretreatment with CytoD did not alter clusterin expression when compared to no pretreatment with CytoD, so all experiments were combined and plotted in Figure 2f. Cells treated with myelin were washed once with PBS prior to RNA preparation.

### Cytokine and H_2_0_2_ *in vitro* treatment

OPCs were treated with 10ng/ml TNFα, 10ng/ml IFNγ, 10µM H_2_0_2_ or the relevant control for 3 hours and then processed for qPCR.

### ELISA

OPCs were grown as described above. A 10cm dish of approximately 2 million OPCs was treated with 3µM Aβ or an equivalent volume of vehicle for 72 hours. Cells were lysed for 15 minutes in RIPA buffer (PBS, 1% Triton X-100, 0.5% deoxycholic acid, 1% sodium dodecyl sulfate) supplemented with 1x protease inhibitor (MedChem Express HY-K0010). The insoluble material was removed by spinning the lysate at 15,000g for 15 minutes. The resulting lysate was used to quantify the amount of clusterin present in each sample using the Mouse Clusterin ELISA Kit (Thermo Fisher EM18RB) according to manufacturer’s instructions.

### Luminex Assay

OPCs were grown as described above. Cells were treated with 8µg/ml clusterin for 24 hours (1 replicate) or 72 hours (1 replicate). Media was then collected and spun to remove all cellular debris. Media was concentrated using a 3KD cutoff concentrator column (Thermo Fisher 88526).

The resulting concentrate was analyzed using the Milliplex 32-plex Mouse Cytokine/Chemokine Panel (Millipore Sigma MCYTMAG-70K-Px32) according to manufacturer’s instructions.

### Apoptotic cell preparation and *in vitro* treatment

To create apoptotic cells, Jurkats were treated with 150mJ of UV energy and incubated for 2-4 hours at 37°C in complete media consisting of RPMI 1640 Media (Gibco, 11875101), 10% FBS (R&D Systems, S12450H), and Pen-Strep (Gibco, 15140122). Apoptotic cells were washed and added to cultured OPCs at approximately a 1:1 ratio for 6 hours. OPCs were washed once prior to RNA isolation.

### Aβ Injections

C57BL/6J mice (8-14 weeks) were injected with CypHer-Aβ (ipsilateral) and NHS-Fluorescein (contralateral, Thermo Fisher 46410) as previously described^60^. Briefly, mice were anesthetized with a mixture of ketamine and xylazine and a small burr hole was drilled in the skull. 1µL of 100µM Aβ was injected at a speed of 200nL/minute into the right hemisphere. 1µL of 200µM NHS-Fluorescein diluted in PBS was injected into the left hemisphere at the same speed. Injections were targeted for 2mM lateral, 0mM anterior, and -1.5mM deep relative to bregma. Mice were given ketoprofen following surgery and were euthanized 12 hours post-injection.

### Immunofluorescence

Mice were deeply anesthetized with pentobarbitol and subsequently perfused with 5 units/mL heparin in saline followed by 10% buffered formalin, each for approximately one minute. For brain tissue, brains were rapidly dissected and post-fixed in 10% buffered formalin overnight at 4°C. Tissue was then transferred into 30% sucrose in PBS and allowed to sink for at least 24 hours. Brains were frozen in OCT, sectioned, and stored in PBS plus 0.02% NaAz until further staining.

Tissue or cultured cells were blocked with PBS, 1% BSA, 0.5% Triton-X 100, 2% normal donkey serum, and 1:200 CD16/CD32 (14-0161-82, 1:200, eBioscience) for at least one hour at room temperature. For stains utilizing a mouse primary antibody, tissue was blocked in Mouse on Mouse Blocking Reagent (MKB-2213, Vector Laboratories) according to manufacturer’s instructions for at least 1 hour at room temperature. Samples were incubated in primary antibodies overnight at 4°C with gentle agitation. Samples were then washed three times in TBS containing 0.3% Triton-X 100 and incubated in secondary antibodies overnight at 4°C with gentle agitation. Following secondary incubation, samples were stained with Hoechst (1:700, ThermoFisher Scientific, H3570) for 10 minutes at room temperature, washed three times in TBS containing 0.3% Triton-X 100, and mounted on slides using Aqua Mount Slide Mounting Media (Lerner Laboratories). Images were collected on a Leica TCS SP8 confocal microscope and processed using Fiji.

### Antibodies for Immunofluorescence

Primary antibodies used for immunofluorescence were PDGFRα (1:200, R&D Systems, AF1062), Olig2 (1:200, Millipore, MABN50), GFP-488 (1:400, Fisher Scientific, A21311), GFP (1:1000, Invitrogen, A10262), Clusterin (1:250, Abcam, AB184100), and Aβ (1:300, Cell Signaling Technology, 8243S). Secondary antibodies used were Donkey anti-Goat Cy3 (2μg/mL, Jackson ImmunoResearch, 705-165-147), Donkey anti-Mouse 647 (2μg/mL, Jackson ImmunoResearch, 715-605-150), Donkey anti-Mouse 546 (2μg/mL, Life Technologies, A10036), Donkey anti-Chicken 488 (2μg/mL, Jackson ImmunoResearch, 703-545-155), Donkey anti-Goat 488 (2µg/ml, Jackson ImmunoResearch, 705-545-147), Donkey anti-Rabbit Cy3 (2µg/ml, Jackson ImmunoResearch, 711-165-152), Donkey anti-Rabbit 647 (2µg/ml, Jackson ImmunoResearch, 711-605-152), and Donkey anti-Goat 647 (2μg/mL, Invitrogen, A21447).

### Isolation of CNS cells

To prepare cells for single-cell sequencing, adult mice (8-20 weeks) were anesthetized with pentobarbitol and subsequently perfused with 5 units/mL heparin in saline for approximately one minute. Brains were rapidly dissected and finely minced. For single-cell sequencing experiments, tissue was digested in HBSS with calcium and magnesium (Gibco, 14025-092) supplemented with 20 units per mL papain (Worthington Biochemical LS003126) and 50 units per mL DNase (Worthington Biochemical, LS002139). Tissue was digested at 37°C with gentle shaking for 45 minutes, with trituration after every 15-minute interval to dissociate the tissue. Following digestion, a 40% Percoll gradient (GE Healthcare, 17-0891-01) was used to remove myelin and other debris from the samples. Resulting single-cell suspensions from 4-5 mice were pooled for each sequencing sample and subsequently stained for FACS sorting.

### FACS sorting

For single-cell sequencing experiments, single-cell suspensions were stained for 30 minutes at room temperature with the following antibodies: O4-APC (O4, 10µL/test, Miltenyi, 130-095-891), CD11b-e450 (M1/70, 0.5 µL/test, eBioscience, 48-0112-82), TER119-APC/Cy7 (TER-119, 1.25 µL/test, Biolegend, 116223), PDGFRα-PE/Cy7 (APA5, 0.625 µL/test, Invitrogen, 25-1401-82), CD45-PerCP/Cy5.5 (30-F11, 0.5 µL/test, eBioscience, 45-0451-82), and CD16/31 (93, 0.5 µL/test, Invitrogen, 14-0161-82). Viability was determined using Ghost Dye Violet 510 (0.5 µL/test, Tonbo biosciences, 13-0870). Cells were sorted using a 16-color BD influx cell sorter. Cells used for sequencing were gated on live/singlets/TER119-/CD45-/CD11b-/YFP+. Following sorting, cells were washed three times with 0.04% BSA and then processed for sequencing according to the 10x Genomics protocol.

### Flow Cytometry

OPCs were incubated with 3µM CypHer-Aβ with 8µg/ml clusterin, 1µM CytoD, or a PBS vehicle control for 90 minutes. Cells were removed from the plate with 0.25% Trypsin-EDTA (Gibco 25200056), washed, and stained with Ghost Dye Violet 510 (0.5 µL/test, Tonbo biosciences, 13-0870). Cells were analyzed using a 3 laser, 10-color Gallios flow cytometer (Beckman-Coulter).

### Single-Cell Sequencing and Analysis

#### Library Preparation and Sequencing

Samples were processed for single-cell sequencing according to manufacturer’s instructions using the Chromium Next GEM Single-cell 3’ Reagent Kit (10xGenomics) and Chromium Controller (10xGenomics). Single-cell libraries were sequenced using the NextSeq 500 Sequencing System (Illumina). Library preparation and sequencing was completed by the Genome Analysis and Technology Core at the University of Virginia.

#### Quantification

All steps of the quantification process were performed with Cellranger. The fastq files for the samples were quantified using the mkfastq utility and were quantified against the mm10 mouse genome with the count utility.

#### Pre-processing

Seurat was used for the single-cell analysis^61, 62^, and for each of the sequencing run datasets, we followed the same procedure. First, a QC step was performed to identify and remove cells that were potential outliers. This included removing potential multiplets (i.e., cells that had clear outlier gene expression) and cells that had approximately ten percent or more of mitochondrial gene expression (i.e., cells that were likely to have high technical variation). After filtering out these suspect cells, the data was normalized and log-transformed (using the ’LogNormalize’ method), unwanted sources of technical variation were regressed out (i.e., the number of detected molecules and mitochondrial contribution to gene expression) ^63^, and the counts were scaled.

#### Integration

To make comparative analyses possible between multiple sequencing run datasets, the datasets were integrated with Seurat using the alignment strategy described previously^61^. The first step was to select the genes to be used as the basis for the alignment. The union of the 1000 genes with highest variability in each of the datasets was taken and then filtered this down to only those genes found in each of the datasets, resulting in 2,285 genes for the alignment. Next, the common sources of variation between the six datasets (3 sequencing runs with 2 samples each) was identified by running a canonical correlation analysis (CCA) with the highly variable genes as features. By examining the strength of the thirty calculated canonical correlations (CCs), the first twelve CCs were identified to be driving the variation between the datasets. The subspaces ^61^ (i.e., the first twelve CCs) were then aligned, resulting in an integrated dataset with features on a common scale.

#### Analysis

Seurat was used on the aligned dataset to identify eight clusters of cells, and then t-SNE was used to visualize the similarity between cells. Next, cell types were assigned to these clusters based upon the expression of pre-defined marker genes, and then identified cluster markers by finding the differentially expressed genes in one cluster compared to all other clusters (one-vs-all). All analyzed single-cell sequencing data has been uploaded in a searchable database located at http://165.22.7.10:3838/seurat_viewer/seurat_viewer_4.Rmd. The sequencing data has been

### Subclustering

Reclustering was performed in Seurat (version 3). Briefly, cells from each original cluster of interest were used to create new Seurat objects consisting of only cells from the clusters of interest. Next, the count data was split by sex and re-normalized with the command: “NormalizeData(object, verbose = FALSE, normalization.method = “LogNormalize”, scale.factor = 10000, assay = “RNA”)”. The Seurat function “FindVariableFeatures” with the “vst” method was used to identify the 2,000 most variable genes in each sex. Next, FindIntegrationAnchors was applied with dims = 1:30 to integrate the data with CCA using the first thirty CC vectors. “IntegrateData” was called to integrate the two sexes. Next, the call: “ScaleData(object, scale.max=10, model.use=“linear”, use.umi=FALSE, do.scale=TRUE, do.center=TRUE, block.size=1000, min.cells.to.block=3000,verbose = FALSE)” was used to re-center and scale the counts matrix for each new Seurat object representing a cluster. “RunPCA(object, npcs = 30, verbose = FALSE)” was used to perform PCA, and “RunTSNE(object,dims.use = 1:14, max_iter=2000)”, “FindNeighbors(object, reduction = “pca”, dims = 1:14)”, and “FindClusters(object, resolution = 0.6)” were called to identify new sub-clusters. Visualizations were performed using the Seurat function “VlnPlot”.

### Multiplex RNAscope (Mouse)

C57B/6J mice (8-10 weeks) from Jackson were anesthetized with pentobarbitol and subsequently perfused with ice-cold 5 units/mL heparin in saline for approximately 1 minute. Brains were rapidly dissected, flash frozen in OCT (Fisher Healthcare, 4585), and stored at -80°C until further processing. Frozen tissue was cut sagittally (15μm), immediately slide-mounted, allowed to dry for approximately one hour at -20°C and then stored at -80°C. All tissue was used within three months of dissection.

Tissue was processed using the V1 RNAscope Fluorescent Multiplex Reagent Kit (Advanced Cell Diagnostics, 320850) according to manufacturer’s instructions. Briefly, tissue was fixed for 15 minutes in 10% buffered formalin (Fisher Scientific, 23-245685) at 4°C, dehydrated, and then incubated in Protease IV (Advanced Cell Diagnostics, 320850) at room temperature for 30 minutes. Target probes were hybridized to the tissue for two hours at 40°C, followed by hybridization of AMP1-FL (30 minutes, 40°C), AMP2-FL (15 minutes, 40°C), AMP3-FL (30 minutes, 40°C), and AMP4-FL (15 minutes, 40°C). Samples were counterstained with supplied DAPI or Hoechst 33342 (1:700, ThermoFisher Scientific, H3570) and mounted on slides using ProLong Glass Antifade Mountant (ThermoFisher, P36980). The following target probes were used: *Olig1* (Advanced Cell Diagnostics, 480651-C2), *Olig2* (Advanced Cell Diagnostics, 447091-C2), *Pdgfrα* (Advanced Cell Diagnostics, 480661), Clusterin (Advanced Cell Diagnostics, 427891-C3), *Gpr17* (Advanced Cell Diagnostics,318131-C3), RNAscope 3-plex Positive Control Probes (Advanced Cell Diagnostics, 320881), and RNAscope 3-plex Negative Control Probes (Advanced Cell Diagnostics, 320871). Sections were imaged using a Leica TCS SP8 confocal microscope.

### RNAscope Quantification

Following imaging, max projected confocal images were analyzed using CellProfiler Software. RNA expression per cell was quantified using a modified version of a previously published pipeline^64^. Briefly, automated steps were used to draw nuclear masks and subsequently quantify the number of RNA puncta from each channel that colocalized with each nuclear mask. Threshold values for each channel were set based on negative control images. Automatic nuclear identification was reviewed and any nuclear mask that clearly contained a large group of nuclei or was located on the edge of an image such that part of the nuclei was not visible was excluded from further analysis. Cells were considered positive for an OPC marker (*Pdgfra*, *Olig1*, or *Olig2*) if four or more puncta colocalized with a particular nucleus to account for background in the assay ^65^. OPCs were defined by the co-expression of two canonical OPC transcripts encoding for cell surface markers (*Pdgfra* or *Cspg4*) and oligolineage transcription factors (*Olig1* or *Olig2*). The number of transcripts of cluster markers clusterin or *Gpr17* were recorded for each identified OPC. OPCs were considered *Clu*- or *GPR17*-if they contained 10 or fewer puncta.

### Multiplex RNAscope (Human)

Human tissue was embedded in paraffin and cut into 15µM sections. Slices were heated for 48 hours to allow attachment. Tissue was subsequently processed using the V2 RNAscope Fluorescent Multiplex Reagent Kit (Advanced Cell Diagnostics, 323100) according to manufacturer’s instructions. Briefly, tissue was dehydrated using xylene and ethanol wash, treated with H_2_0_2_, and then incubated in Target Retrieval Buffer at 95°C for 15 minutes. Tissue was incubated with the supplied Protese Plus regent for 30 minutes. Target probes were hybridized to the tissue for two hours at 40°C, followed by hybridization of AMP1-FL (30 minutes, 40°C), AMP2-FL (15 minutes, 40°C), AMP3-FL (30 minutes, 40°C), and AMP4-FL (15 minutes, 40°C). Tissue was then treated with an HRP reagent for a single probe, followed by a unique secondary, and an HRP blocker. This process was repeated for each probe used. Samples were counterstained with supplied DAPI or Hoechst 33342 (1:700, ThermoFisher Scientific, H3570) and mounted on slides using ProLong Glass Antifade Mountant (ThermoFisher, P36980). The following target probes were used: Human *OLIG2* (Advanced Cell Diagnostics, 424191-C2), Human *PDGFRA* (Advanced Cell Diagnostics, 604481), Human *CLU* (Advanced Cell Diagnostics, 584771-C3), and the RNAscope 4-plex Negative Control Probes (Advanced Cell Diagnostics, 321831). The following dyes were used: Opal 520 (Akoya Biosciences, FP1487001), Opal 690 (Akoya Biosciences, FP1497001), and Opal 620 (Akoya Biosciences, FP1495001). Sections were imaged using a Leica TCS SP8 confocal microscope.

### Immunohistochemistry (Human)

5 μm thick tissue sections of formalin fixed, paraffin embedded (FFPE) brain tissue specimens were rehydrated in Xylene and descending alcohol series and heat-induced epitope retrieval (HIER) was performed in a pressure cooker (Tintoretriever, Bio SB) for 15 min at high pressure in a 0.05% Tween-20 solution. Endogenous peroxidase was quenched by incubation of sections in 1.5% hydrogen peroxide/0.005% Triton-X-100 diluted in pH 7.4 sterile phosphate buffered saline (PBS) (Invitrogen) for 20 min, following multiple washes in tap water and subsequently, 0.1 M Tris, pH 7.6. Non-specific antibody binding was minimized with sections incubated in 2% FBS/0.1 M Tris, pH 7.6. Clusterin (Proteintech) primary antibody was diluted in 2% FBS/0.1 M Tris, pH 7.6 at a dilution of 1:500. Sections were incubated with primary antibody over night at 4°C, washed one time in 0.1 M Tris, pH 7.6, followed by 2% FBS/0.1 M Tris, pH 7.6 for 5 min, incubated in goat ant-rabbit IgG HRP Conjugated secondary antibody (Millipore Sigma) for 1 hour, additionally washed one time in 0.1 M Tris, pH 7.6, followed by 2% FBS/0.1 M Tris, pH 7.6 for 5 min, and incubated in VectaStain ABC Peroxidase HRP Kit (diluted in 2% FBS/0.1 M Tris, pH 7.6 at 1:1000) for 1 hour. After a final wash in 0.1 M Tris, pH 7.6 for 5 min, immunocomplexes were visualized using the Vector Laboratories ImmPACT DAB Peroxidase (HRP) 3,3′-diaminobenzidine. Tissue sections were counterstained with hematoxylin (Sigma Aldrich, St. Louis, MO) for 2 minutes, dehydrated in ascending alcohol series and Xylene and cover slipped using Cytoseal 60 mounting medium (Thermo Scientific). For analysis of stains, slides of frontal cortex specimens stained with Clusterin antibodies were scanned on an Aperio AT2 scanner (Leica biosystems) at 20x magnification and digital slides analyzed using the QuPath platform (version 0.3.1, https://QuPath.github.io/ , PMID: 29203879) on a Dell PC (Intel® Xeon® W-1270 CPU @ 3.40GHz/ 64 GB RAM/ 1 TB SSD Hard Drive) running Windows 10. Cortex and white matter were annotated for regional analysis. After exclusion of tissue and staining artifacts we used the ‘Positive Pixel Detection’ tool (Downsample factor 2, Gaussian sigma 1 μm, Hematoxylin threshold (‘Negative’) 1.5 OD units, DAB threshold (‘Positive’) 0.5 OD units) to determine the percentage of area covered by Clusterin staining.

### Combined in situ Hybridization and Immunohistochemistry (Human)

For in-situ hybridization, 5µm thick paraffin-embedded tissue sections on slides were rehydrated in xylene and series of ethanol solutions (100%, 90%, and 70%). Following air drying for 5 min at RT, slides were incubated with RNAscope® Hydrogen peroxide for 10 min at RT, followed by 3 washing steps in distilled water. Antigen retrieval was performed in a steam bath for 15 min using RNAscope® 1x target retrieval reagent. After a rinse in distilled water and incubation in 10% ethanol for 3 min slides were air dried at 60°C. Subsequently, slides were incubated with RNAscope® Protease plus reagent for 30 min at 40°C in a HybEZ™ oven, followed by 3 washes in distilled water. Slides were then incubated with the following RNAscope® probe for 2 hours at 40°C in a HybEZ™ oven: Hs-Clusterin (cat. number 606241). Following washes with 1X Wash buffer, slides were incubated with RNAscope®AMP1 solution for 30 min at 40°C. Subsequent incubations with other RNAscope® AMP solutions, followed each by two washes with 1x RNAscope® Wash buffer were completed as follows: AMP2 – 15 min at 40°C; AMP3 – 30 min at 40°C; AMP4 – 15 min at 40°C; AMP5 – 30 min at RT and AMP6 – 15 min at RT. After two washes in 1x RNAscope® wash buffer slides were incubated in RNAscope® Fast RED-B and RED-A mixture (1:60 ratio) for 10 min at RT, followed by two washes in tap water. For in situ hybridization/immunohistochemistry double labeling, sections were incubated in 2% FBS/0.1 M Tris, pH 7.6 for 5 min following RNAscope® Fast RED incubation and two washes in tap water. Primary antibodies were diluted in 2% FBS/0.1 M Tris, pH 7.6 at the following dilutions: Ab5 (PMID: 16341263), 1:1000. Sections were incubated with primary antibody over night at 4°C, washed two times in 0.1 M Tris, pH 7.6 for 5 min each and incubated with biotinylated secondary antibody (Vector Laboratories; Burlingame, CA) diluted in 2% FBS/0.1 M Tris, pH 7.6 for 1 hour at room temperature. An avidin-biotin complex (ABC) system (Vectastain ABC Elite kit; Vector Laboratories, Burlingame, CA) was used to enhance detection of the immunocomplexes, which were visualized using the chromogen 3,3′-diaminobenzidine (DAB kit; KPL, Gaithersburg, MD). Tissue sections were counterstained with hematoxylin (Sigma Aldrich, St. Louis, MO), air dried at 60°C for 30 min and cover slipped using EcoMount™ mounting medium (Biocare Medical).

### RT-qPCR

RNA was extracted from samples using the ISOLATE II RNA Micro Kit (Bioline, BIO-52075) or the ISOLATE II RNA Mini Kit (Bioline, BIO-52073). Isolated RNA was reverse transcribed using the SensiFAST cDNA Synthesis Kit (Bioline, BIO-65054) or the iScript cDNA Synthesis Kit (Bio-Rad, 1708891). RT-qPCR was performed using the SensiFAST Probe No-ROX Kit (Bioline, BIO-86005) and TaqMan probes for *Gapdh* (ThermoFisher, Mm99999915_g1), *Mbp* (ThermoFisher, Mm01266402_m1 and Mm01266403_m1), *Plp1* (ThermoFisher, Mm00456892_m1), *Cnp* (ThermoFisher, Mm01306641_m1), *Myrf* (ThermoFisher, Mm01194959_m1), and *Clu* (ThermoFisher, Mm00442773_m1). Additionally, the SensiFAST SYBR No-ROX Kit (Bioline, BIO-98005) was used with primers for *Plp1* (Forward: GCCCCTACCAGACATCTAGC, Reverse: AGTCAGCCGCAAAACAGACT) and *Myrf* (CGGCGTCTCGACAGCCTCAA, Reverse: GACACGGCAAGAGAGCCGTCA). Data was collected using the CFX384 Real-Time System (Bio-rad).

### RNA Flow Cytometry

Isolated cells were stained according to the PrimeFlow RNA Assay kit (ThermoFisher Scientific, 88-18005-204). Probes used include *Olig2* (Affymetrix, VPFVKKV-210), *Gpr17* (Affymetrix, VPGZE6T-210), *Clu*(Affymetrix, VB10-3283998-210) and *Pdgfra* (Affymetrix, VB6-3197712-210). A probe targeting *Actb* (Affymetrix, VB1-10350-210) was used as a positive control to ensure good RNA quality. Samples were run using a 16-color Life Technologies Attune Nxt flow cytometer and data was analyzed using FlowJo software.

### Statistical Analysis

Statistical analysis of all data (except single-cell sequencing data) was done using Prism 9 (Graphpad software). Significance was set at p<0.05.

---

**Extended Data Figure 1:**
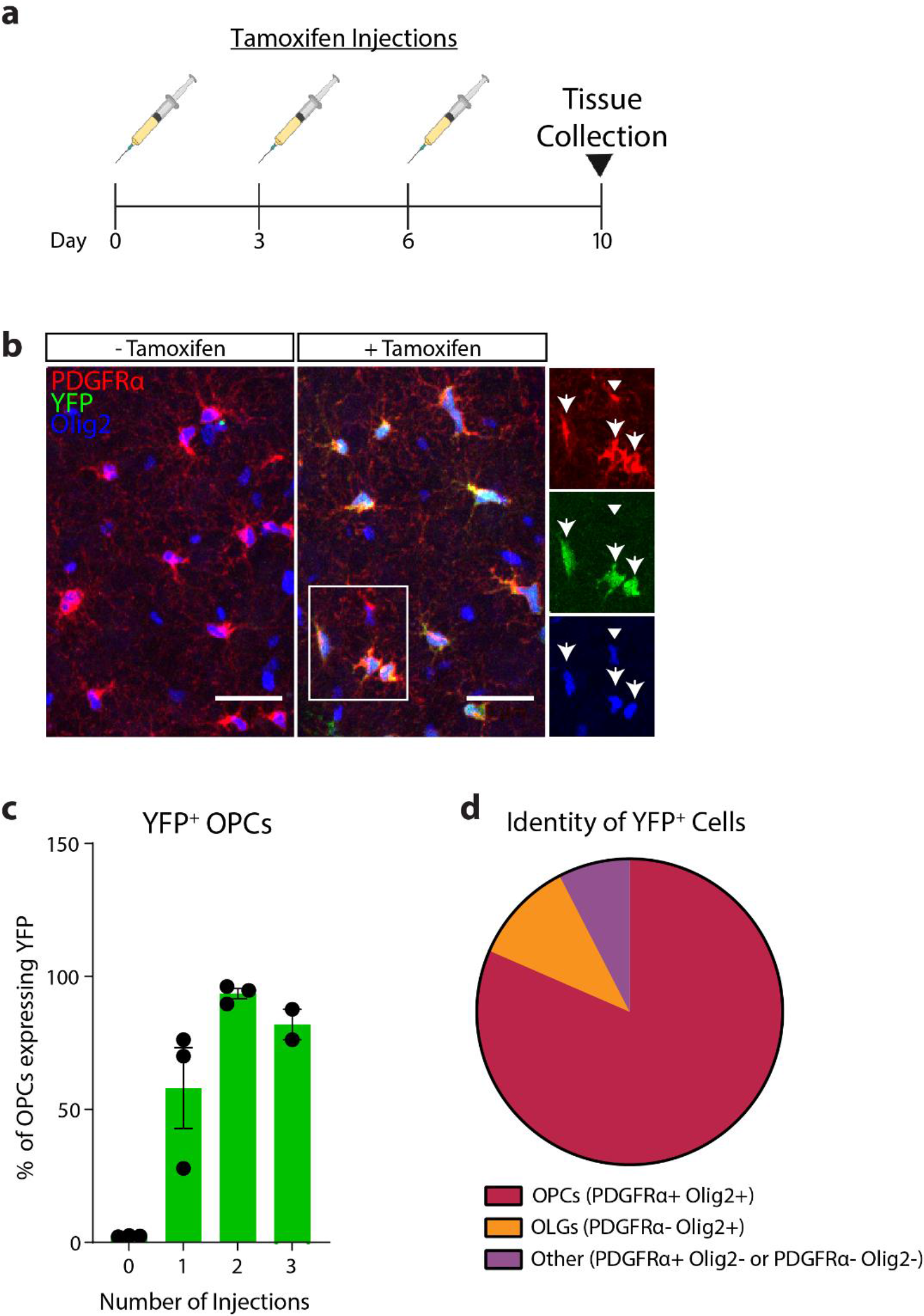
Validation of *PDGFRα-CreER*; *R26-EYFP* reporter mouse. **a**, Timeline of tamoxifen injections and tissue harvest used to validate YFP expression in OPCs and titrate optimal tamoxifen dosing paradigm. **b**, Immunofluorescence of PDGFRα (red), Olig2 (blue), and YFP (green) in *PDGFRα-CreER*; *R26-eYFP* mice receiving no tamoxifen injections (-Tamoxifen) or 2 tamoxifen injections (+Tamoxifen). Arrows represent OPCs expressing YFP and arrowheads represent OPCs lacking YFP expression. Scale bar = 30µM. **c**, Percentage of OPCs (PDGFRα+/Olig2+) that also express YFP following 0, 1, 2, or 3 tamoxifen injections (0 injections *n*=3, 1 injection *n*=3, 2 injections *n*=3, 3 injections *n*=2; from three independent experiments). Error bars represent SEM. **d**, Proportion of YFP+ cells identified as OPCs (PDGFRα+/Olig2+), Oligodendrocytes (OLG, PDGFRα-/Olig2+), or neither of these cell types (Other, PDGFRα+/Olig2- or PDGFRα-/Olig2-) following 2 tamoxifen injections (*n*=3; from two independent experiments). Data quantified in **c** and **d** include images from the prefrontal cortex, hippocampus, and corpus callosum.

**Extended Data Figure 2:**
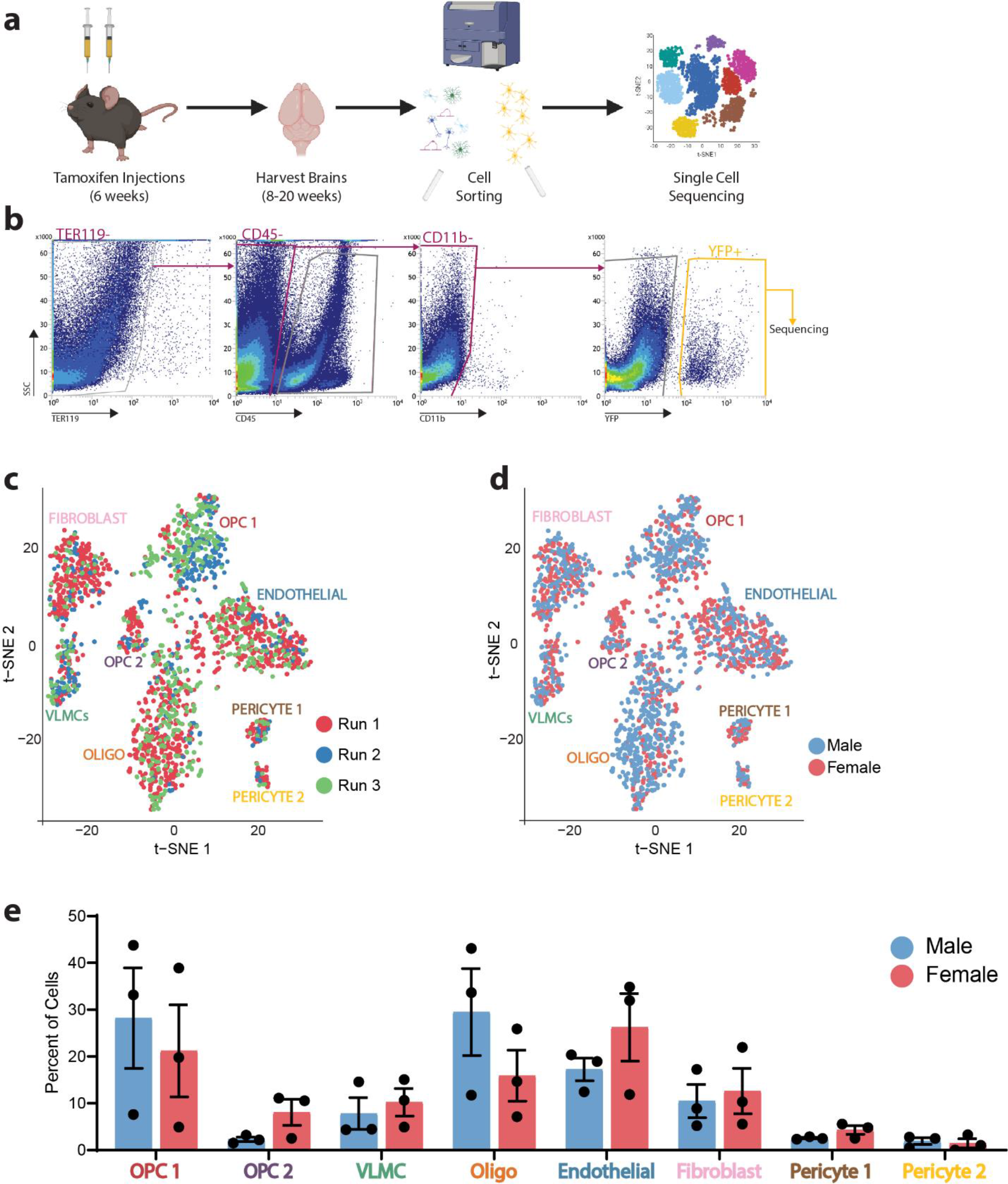
Isolation of YFP+ cells from *PDGFRα-CreER*; *R26-EYFP* brains for single-cell sequencing. **a**, Experimental strategy used for the isolation and single-cell sequencing of cells analyzed in Figure 1f. **b**, Gating strategy for YFP+ cell sorting following Live/Scatter/Singlet gating. **c**, tSNE map depicting cell clusters colored by sequencing run. **d**, tSNE map depicting cell clusters colored by sex. **e**, Percent of each sequencing sample belonging to each sample (Male *n*=3, Female *n*=3; from three independent experiments). No statistically significant differences in cell abundance were found in any cluster between males and females using unpaired t-tests.

**Extended Data Figure 3:**
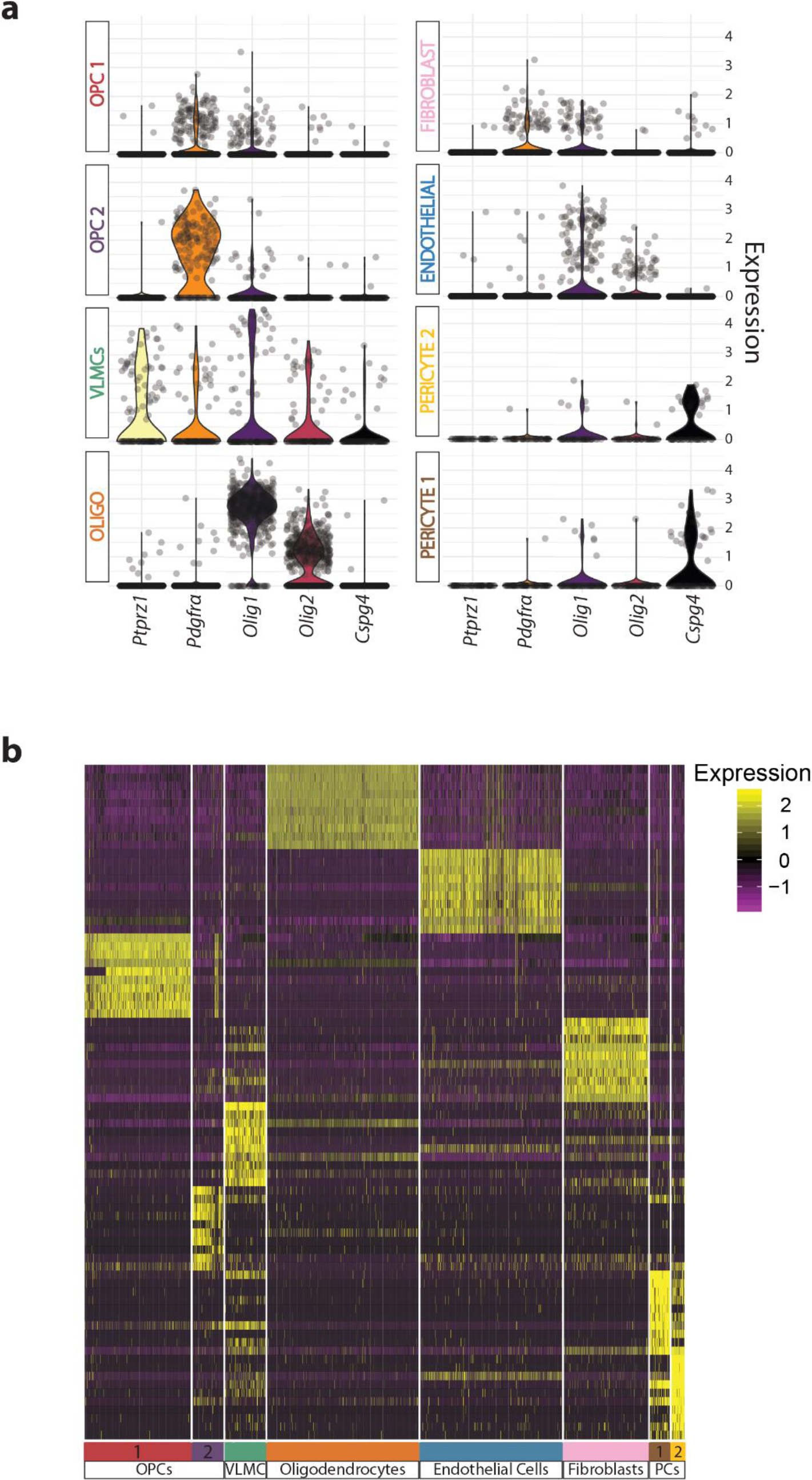
Gene expression of single-cell sequencing clusters. **a**, Violin plots depicting expression of common OPC markers in each cluster. Each dot represents a cell. Expression value is plotted on the y-axis. **b**, Heatmap depicting the scaled and log-normalized expression values of the top 10 most highly enriched genes in each cluster. PC=Pericytes.

**Extended Data Figure 4:**
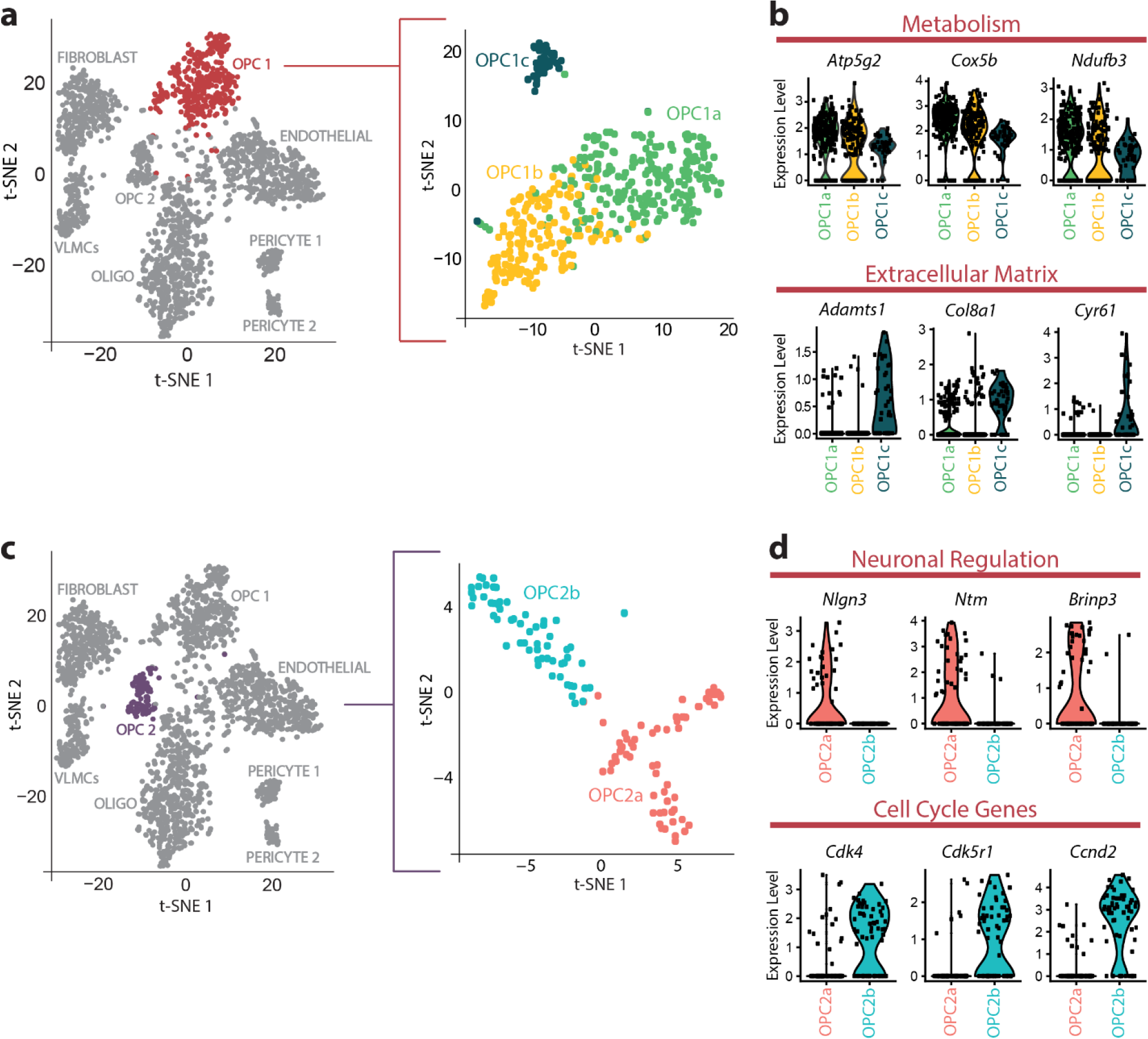
Subclustering reveals potential functions of OPC clusters. **a**, Subclustering of OPC1 into 3 subclusters (OPC1a, OPC1b, and OPC1c). **b**, Violin plots depicting expression of genes related to metabolism that are significantly upregulated in OPC1a and genes related to the extracelluar matrix that are significantly upregulated in OPC1c. Each dot represents a cell. **c**, Subclustering of OPC2 into 2 subclusters (OPC2a and OPC2b). **d**, Violin plots depicting expression of genes related to neuronal regulation that are significantly upregulated in OPC2a and genes related to the cell cycle that are significantly upregulated in OPC2b. Each dot represents a cell.

**Extended Data Figure 5:**
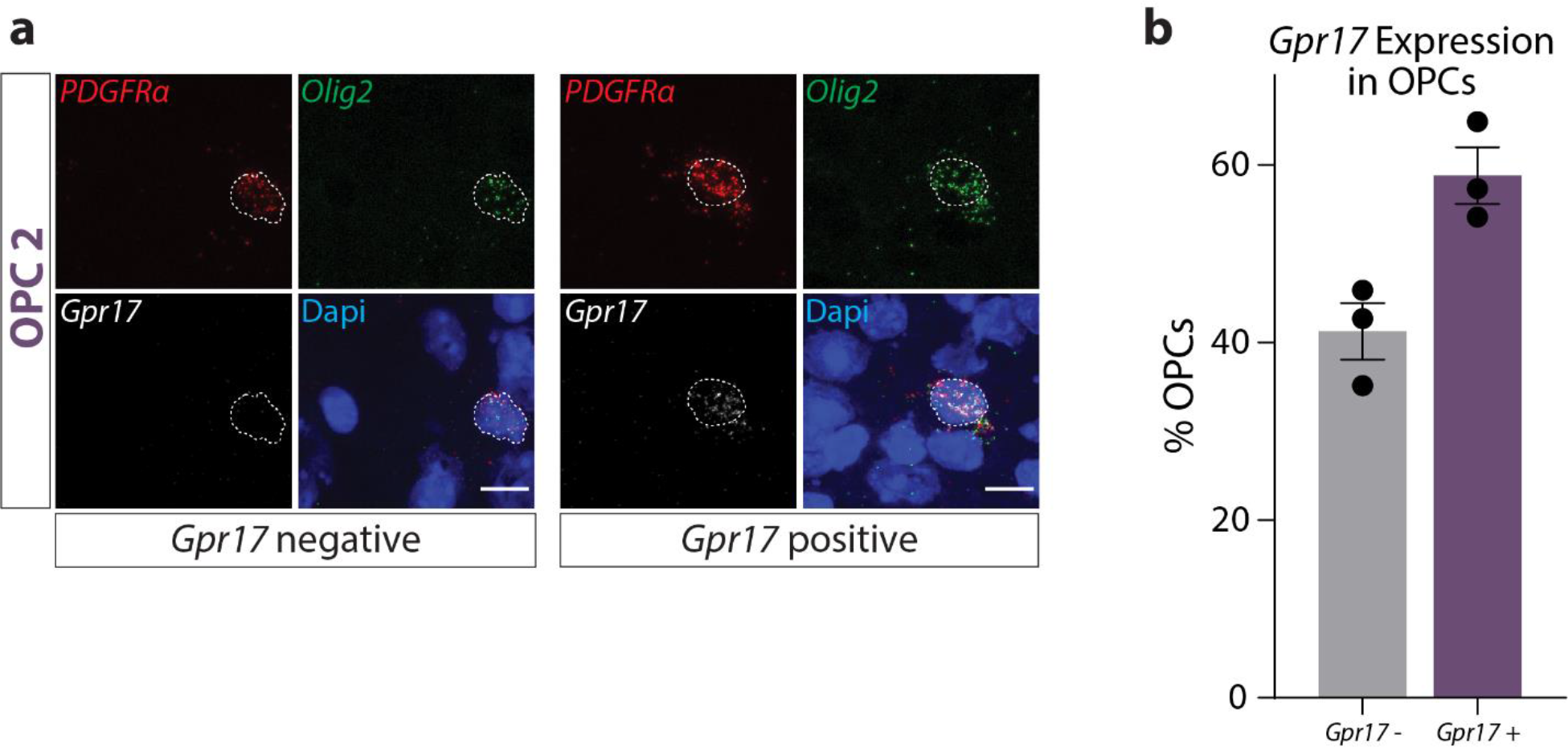
*Gpr17* is expressed by a subset of OPCs *in vivo*. **a**, Representative images of *in situ hybridization* for OPCs (*Pdgfra* in red, *Olig2* in green) expressing or lacking *Gpr17* (white). Scale bar=10µm. **b**, Quantification of *GPR17*+ and *GPR17-*OPCs (depicted in **a**; *n*=3 with 204 total cells analyzed; from two independent experiments). Each sample includes quantification of OPCs from the cortex, hippocampus, corpus callosum, and cerebellum.

**Extended Data Figure 6:**
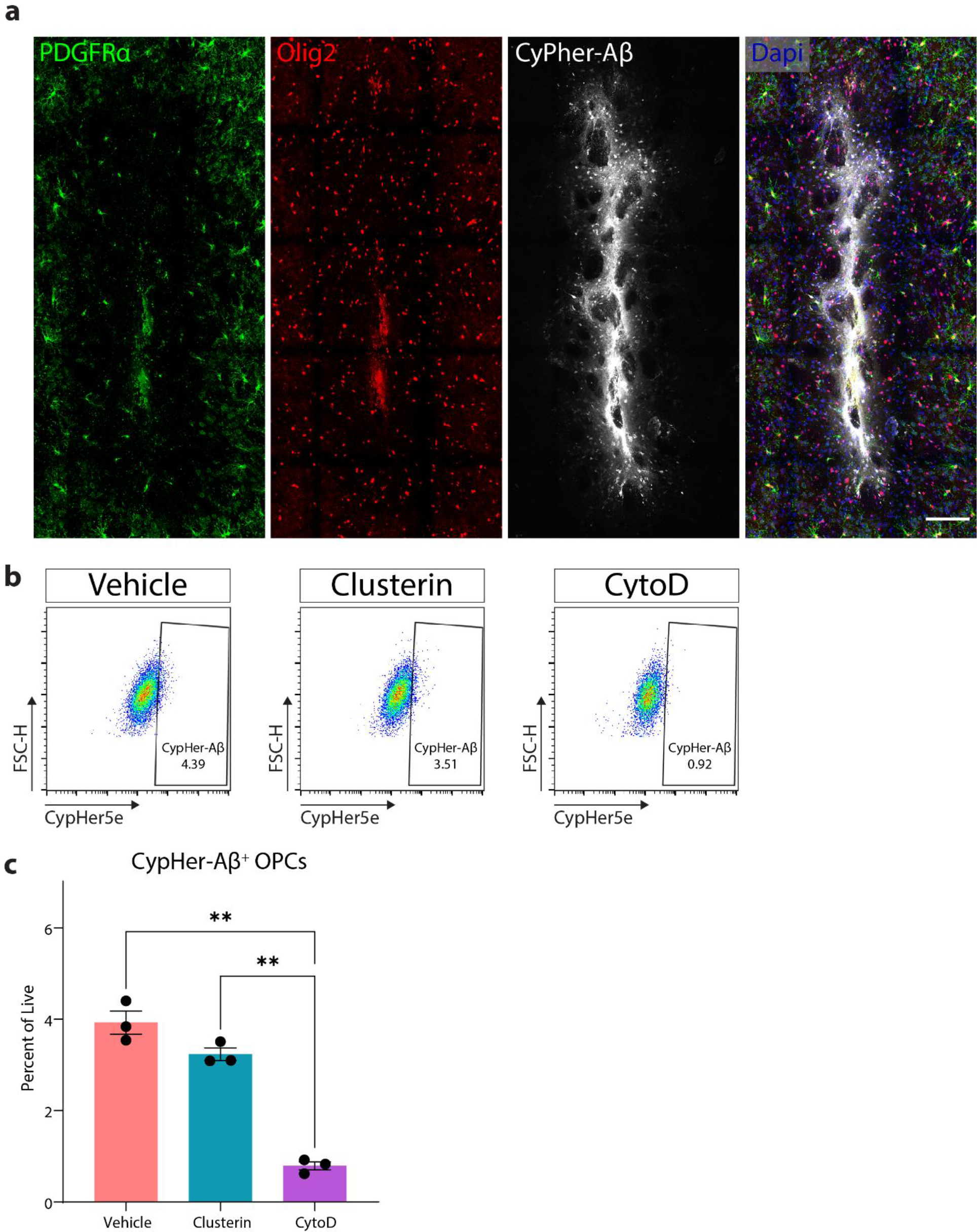
Clusterin does not alter OPC phagocytosis of Aβ. **a**, Low magnification image of OPCs (PDGFRα in green, Olig2 in red) surrounding CypHer5e-labeled Aβ oligomers 12 hours after they were injected into the parenchyma (representative image of *n*=6; from one independent experiment). **b**, Representative flow gating (following singlets/singlets/live gates) of OPCs incubated for 90 minutes with 3µm CypHer5e-labeled Aβ oligomers (all conditions) with the addition of 8µg/ml clusterin or 1µm CytoD (Vehicle *n*=3, Clusterin *n*=3, CytoD *n*=3; from one independent experiment). CypHer+ gate was drawn so that less than 1% of cells in the CytoD samples fell within the positive gate. **c**, Quantification of OPCs staining positive for CypHer-Aβ as depicted in **b**. Statistics calculated using a repeated measures one-way ANOVA with a Tukey’s post-hoc analysis; F(2,4)= 293.3. **p<0.01. All error bars represent SEM.

**Extended Data Figure 7:**
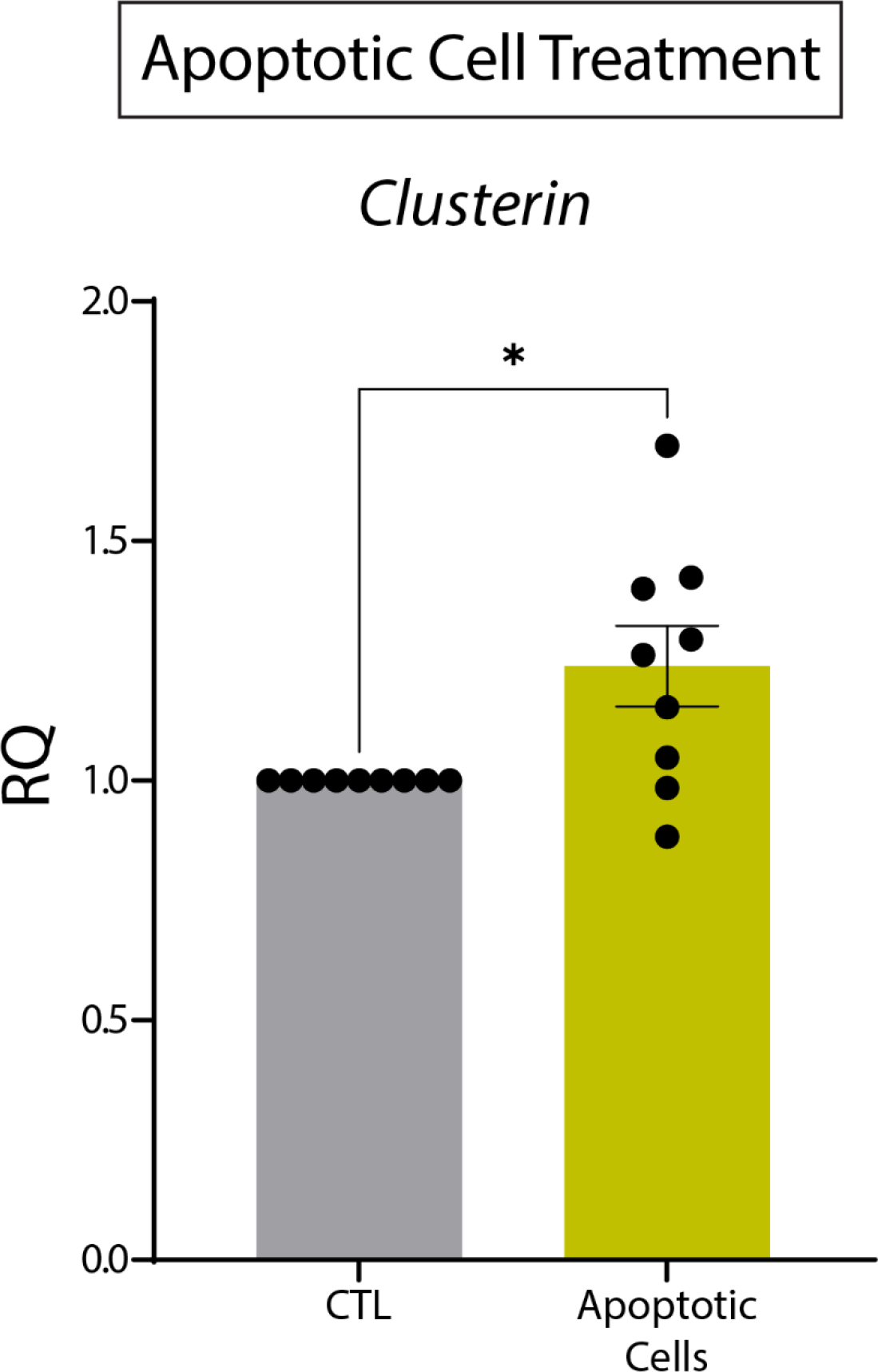
Apoptotic cells drive clusterin expression in OPCs. qPCR analysis of clusterin expression in OPCs following a 6-hour *in vitro* treatment with apoptotic cells (CTL *n*=9, Apoptotic cells *n*=9; from two independent experiments). Statistics calculated using a paired Student’s t-test; t(8)=2.835. *p<0.05. All error bars represent SEM.

**Extended Data Figure 8:**
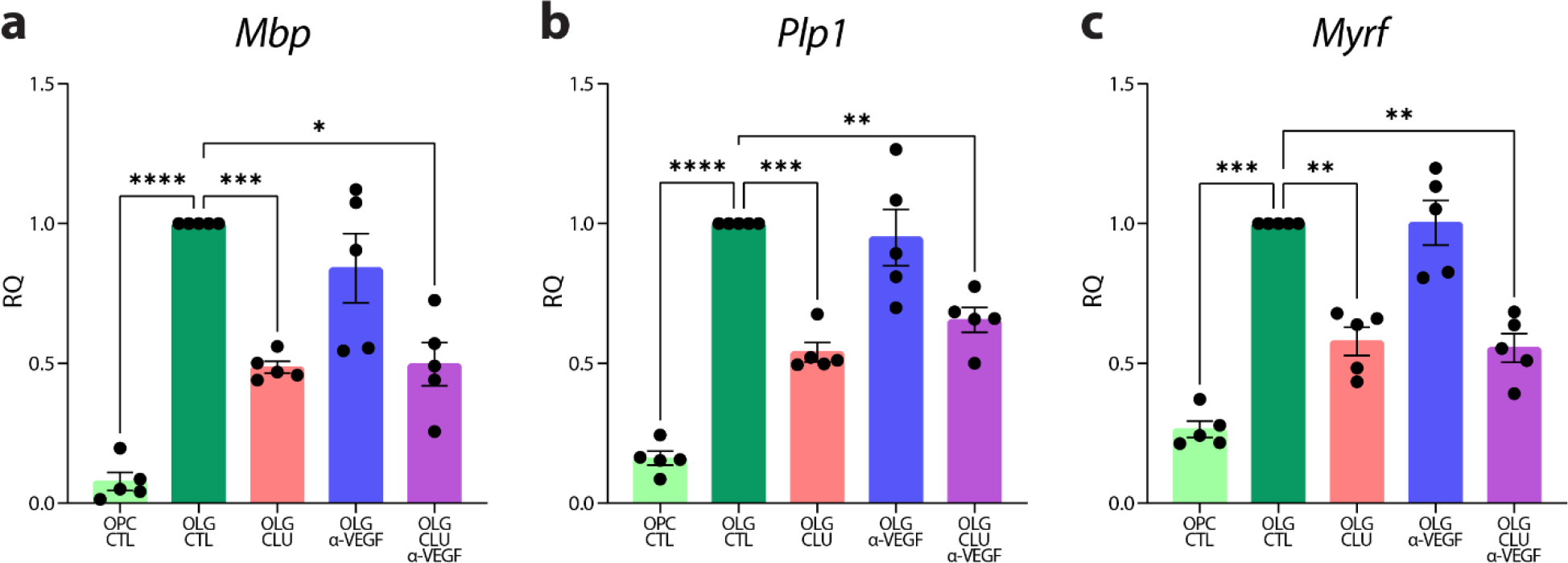
VEGF does not mediate the effect of clusterin on OPC differentiation. Expression of *Mbp* (**a**), *Plp1* (**b**), and *Myrf* (**c**), measured by qPCR in OPCs cultured in proliferation media (OPC CTL), differentiation media (OLG CTL), differentiation media supplemented with 8µg/ml of clusterin (OLG CLU), differentiation media supplemented with 10µg/ml VEGF function-blocking antibody (OLG α-VEGF), or differentiation media supplemented with 8µg/ml of clusterin and 10µg/ml VEGF function-blocking antibody (OLG CLU α-VEGF) for 72 hours (*n*=5 for all conditions; from one independent experiment). Statistics calculated using a repeated measures one-way ANOVA with a Tukey’s post-hoc analysis; *Mbp* F(4,16)=28.66, *Plp1* F(4,16)=49.31, *Myrf* F(4,16)=74.58. *p<0.05, **p<0.01, ***p<0.001 ****p<0.0001. All error bars represent SEM.

**Supplementary Table 1:** List of human cases used for Clusterin analysis (Fig 1a, b). Cases were analyzed by a board-certified neuropathologist according to current diagnostic guidelines for Alzheimer’s disease neuropathologic change (ADNC, PMID: 22101365, 22265587), Cerebral amyloid angiopathy (CAA, PMID: 25716356), Aging-related tau astrogliopathy (ARTAG, PMID: 26659578), Limbic-predominant age-related TDP-43 encephalopathy neuropathologic change (LATE NC, PMID: 31039256) and primary age-related tauopathy (PART, PMID: 25348064). Additional abbreviations used in the table: Braak – Braak stage; CVD – Cerebrovascular disease; Thal – Thal phase; CERAD-Consortium to establish a registry for Alzheimer’s disease

| Case number | Main neuropathologic diagnosis | Secondary neuropathologic diagnoses | Thal | Braak | CERAD | Gender | Age |
| --- | --- | --- | --- | --- | --- | --- | --- |
| control 1 | PART, definite, Braak II |  | 0 | II | none | f | 72 |
| control 2 | PART, Braak I | CAA widespread, mild | 0 | I | none | m | 71 |
| control 3 | No significant pathological findings |  | 0 | 0 | none | f | 55 |
| control 4 | CVD | ADNC low; CAA focal, mild | 1 | II | none | f | 90 |
| control 5 | No significant pathological findings | Atherosclerosis (moderate) | 0 | 0 | none | m | 71 |
| control 6 | No significant pathological findings | Atherosclerosis (moderate) | 0 | 0 | none | m | 88 |
| control 7 | PART, definite, Braak I | ARTAG, subependymal | 0 | I | none | m | 77 |
| control 8 | CVD |  | 0 | 0 | none | m | 72 |
| control 9 | PART, definite, Braak I |  | 0 | I | none | f | 73 |
| control 10 | PART, definite, Braak II |  | 0 | II | none | f | 82 |
| control 11 | PART, definite, Braak II | subacute microinfarct corpus callosum | 0 | II | none | f | 90 |
| control 12 | PART, definite, Braak I |  | 0 | I | none | f | 78 |
| control 13 | PART, definite, Braak II | CAA focal, moderate | 0 | II | none | m | 77 |
| control 14 | PART, definite, Braak I | ARTAG, subependymal | 0 | I | none | m | 51 |
| control 15 | PART, definite, Braak I |  | 0 | I | none | m | 52 |
| AD 1 | ADNC high | CAA widespread, moderate | 4 | V | frequent | m | 84 |
| AD 2 | ADNC high | CAA widespread, moderate ; LATE NC stage 1 | 5 | VI | frequent | m | 81 |
| AD 3 | ADNC high | CAA focal, moderate | 4 | V | frequent | f | 85 |
| AD 4 | ADNC high | CAA widespread, moderate to severe | 5 | V | frequent | m | 74 |
| AD 5 | ADNC high | CAA widespread, moderate | 5 | VI | frequent | m | 83 |
| AD 6 | ADNC high | CAA widespread, moderate | 4 | VI | frequent | m | 75 |
| AD 7 | ADNC high | CAA widespread, moderate | 5 | VI | frequent | m | 70 |
| AD 8 | ADNC high | CAA, moderate, widespread | 5 | V | frequent | f | 86 |
| AD 9 | ADNC high | CAA focal, moderate; LATE NC stage 1 | 5 | VI | frequent | m | 74 |
| AD 10 | ADNC high | CAA, widespread, moderate | 5 | V | frequent | f | 64 |
| AD 11 | ADNC high | CAA widespread, moderate | 5 | VI | frequent | m | 59 |
| AD 12 | ADNC high | CAA widespread, moderate | 5 | VI | frequent | m | 83 |
| AD 13 | ADNC high | CAA focal, mild | 5 | V | frequent | m | 78 |
| AD 14 | ADNC high | CAA focal, moderate | 5 | V | frequent | m | 95 |
| AD 15 | ADNC high | CAA widespread, mild to moderate; LATE NC stage 2 | 4 | V | frequent | f | 78 |
| AD 16 | ADNC high | CAA widespread, mild to moderate | 5 | V | frequent | f | 97 |
| AD 17 | ADNC high | CAA widespread, moderate | 5 | V | frequent | f | 78 |
| AD 18 | ADNC high | CAA widespread, mild | 4 | V | frequent | m | 77 |
| AD 19 | ADNC high | CAA widespread, mild to moderate | 5 | VI | frequent | m | 66 |
| AD 20 | ADNC high | CAA widespread, moderate | 5 | VI | frequent | m | 63 |
| AD 21 | ADNC high | CAA focal, moderate | 5 | VI | frequent | f | 86 |
| AD 22 | ADNC high | CAA widespread, mild; LATE NC stage 1 | 5 | V | frequent | f | 82 |
| AD 23 | ADNC high | CAA focal, mild; LATE NC stage 1 | 5 | V | frequent | f | 90 |
| AD 24 | ADNC high | LATE NC stage 2 | 5 | V | frequent | m | 87 |
| AD 25 | ADNC high | CAA focal, mild to moderate | 5 | V | frequent | m | 73 |
| AD 26 | ADNC high | CAA focal, moderate; LATE NC stage 1 | 5 | VI | frequent | f | 84 |

